# Proximity labeling defines the phagosome lumen proteome of murine and primary human macrophages

**DOI:** 10.1101/2024.09.04.611277

**Authors:** Benjamin L. Allsup, Supriya Gharpure, Bryan D. Bryson

## Abstract

Proteomic analyses of the phagosome has significantly improved our understanding of the proteins which contribute to critical phagosome functions such as apoptotic cell clearance and microbial killing. However, previous methods of isolating phagosomes for proteomic analysis have relied on cell fractionation with some intrinsic limitations. Here, we present an alternative and modular proximity-labeling based strategy for mass spectrometry proteomic analysis of the phagosome lumen, termed PhagoID. We optimize proximity labeling in the phagosome and apply PhagoID to immortalized murine macrophages as well as primary human macrophages. Analysis of proteins detected by PhagoID in murine macrophages demonstrate that PhagoID corroborates previous proteomic studies, but also nominates novel proteins with unexpected residence at the phagosome for further study. A direct comparison between the proteins detected by PhagoID between mouse and human macrophages further reveals that human macrophage phagosomes have an increased abundance of proteins involved in the oxidative burst and antigen presentation. Our study develops and benchmarks a new approach to measure the protein composition of the phagosome and validates a subset of these findings, ultimately using PhagoID to grant further insight into the core constituent proteins and species differences at the phagosome lumen.

## Introduction

Professional phagocytes, including monocytes, macrophages, neutrophils, and dendritic cells, coordinate immune system function in infection^1,2^, cancer^3,4^, aging^5^, and beyond. Central to their role as master regulators of health and disease is their ability to phagocytose diverse cargos including external threats (e.g. pathogens^6–9^ and debris^10,11^) as well as those derived from internal threats (e.g. apoptotic cells^1,12^ and biomolecules with pathologic capacity^5,13,14^). In the context of vaccination, professional phagocytes encounter the delivered antigen and adjuvant, process it for antigen presentation, then migrate to local lymph nodes to mount a full immune response^15^. Together, these observations underscore the importance of the phagosome as an immunologic organelle at the center of immune defense against pathogens and cancers, tissue homeostasis, and successful vaccination. Without proper phagosome function, human health is compromised with increased rates of infection, inflammation, and biomechanical defects leading to disease^16,17^.

The phagosome does not exist within a cell as an organelle until a phagocyte comes into contact with an extracellular body and a phagocytic receptor triggers the active spread of phagocyte membrane around the target until it is fully engulfed^1,18^. The nascent phagosome then undergoes a series of fission and fusion events, acquiring proteins and the resultant biochemical properties from early endosomes before eventually fusing with late endosomes and finally lysosomes^1,19,20^. After its cargo is degraded, the phagosome is resolved as it becomes a cargo-less lysosome^21,22^. Because of its highly dynamic and interactive nature, the phagosome does not exist as an independent organelle, but instead as a mosaic of multiple contributing cell compartments.

Several studies have demonstrated that particular proteins and lipids modulate the acidity, degradative capacity, and oxidative potential of the phagosome^23–28^. Further, these studies have demonstrated that the recruitment of these proteins and lipids, and thus the resulting biochemical environment of the phagosome, changes based on cell type and different phagosomal cargos^29–33^. Given the enormous role that the phagosome plays in human health, it is essential that we have a suite of techniques to comprehensively analyze phagosome composition and function in all of the contexts of phagocytosis.

Techniques have been developed to measure one-dimensional properties of the phagosome such as acidity, degradative capacity, or reactive oxygen species (ROS) generation and have granted insight into the biochemical environment of the phagosome induced by different phagosomal cargos, cell types, and cell environments^29,30,34^. Often these studies nominate proteins for targeted followup based on prior knowledge, and distinct phagosomal states are attributed to one or few proteins^25,27^. However, it is likely that distinct phagosomal states are derived from the integrated contributions of many proteins present at the phagosome. To gain a comprehensive understanding of how phagosome function is altered by many proteins in tandem, proteomic techniques to examine phagosome composition are needed.

Proteomic analysis of the phagosome is complicated by its amorphous nature. Some organelles, such as the nucleus or mitochondrion, can be isolated without contamination from other organelles by cell fractionation and differential centrifugation due to their unique density. Clean preparations of these organelles has enabled high-confidence and unbiased proteomic analyses, allowing for discovery of resident proteins and giving significant insight into their function and dysfunction^35–38^. However, the phagosome is not easily isolated by differential centrifugation because the density of a phagosome is determined by its cargo. As a result, many phagosomes containing biological cargo (e.g. bacteria) have densities similar to other lysosomes and endosomes which prevents recovery of all phagosomes without contaminating the phagosome preparation^26,39,40^.

The problem of phagosome density overlap was originally solved in 1969 by Wetzel and Korn where amoeba were allowed to phagocytose latex beads in place of a biological cargo^41^. Latex beads have a high artificial buoyancy allowing for their pure isolation across a single-step sucrose gradient after mechanical disruption of the outer cell membrane^41^. This method was adapted to analyze phagosomes derived from professional phagocytes by Desjardins and colleagues in 1994^42^, and since then more than 20 phagosome proteomic studies have utilized this cell-fractionation approach to analyze latex- or paramagnetic-bead containing phagosomes isolated by differential centrifugation or magnetic isolation, respectively^32,43–45^.

Despite great insight gleaned from these studies, several limitations remain. First, the fractionation approach does not enable a clean preparation of phagosomes containing biological cargos such as bacteria without extensive optimization^7,26,40^. Second, a large amount of cell input is required to isolate a sufficient amount of phagosomes for analysis by mass spectrometry^32,39,44^. This has limited the phagosome proteomic analysis of cells which are only available in modest abundance (e.g. primary human cells)^46,47^.

Proximity labeling is a strategy which enables *in situ* labeling and subsequent proteomic analysis of organelles which can not be cleanly isolated by cell fractionation^48,49^. Specifically, proximity labeling has been applied to characterize the proteomes of suborganellar features of the mitochondrion^50–52^ as well as the lipid droplet^49^, autophagosome lumen^53^, neuronal synaptic cleft^54^, and chlamydia inclusions^55^, among others. Importantly, the proximity labeling reaction is conducted in live cells with membrane-impermeant radicals which maintains the topology and integrity of the organelle and minimizes potential contaminants introduced by cell fractionation^48,49^.

In this study, we develop PhagoID, an approach capable of performing proximity labeling in the phagosome lumen to identify proteins contributing to phagosome function near the phagosomal cargo. We benchmark multiple proximity labeling approaches and identify a low-cost horseradish peroxidase (HRP)-based (HRP-PhagoID) strategy as an optimal strategy for phagosome proximity labeling *in vitro*. We next combine our optimized labeling protocol with TMT-multiplexed quantitative mass spectrometry and apply HRP-PhagoID to the phagosomes of an immortalized murine macrophages and primary human monocyte-derived macrophages, yielding high-confidence proteomes of the phagosome lumen from both species. HRP-PhagoID applied to both murine and human macrophages recovers a phagosome proteome enriched for proteins known to reside in endosomal and lysosomal compartments, consistent with the model of phagosome progression and previously published phagosome proteome datasets. Our datasets also highlight a set of proteins with little or no previous evidence of phagosome residence, and we note that many of these proteins are annotated as residing in diverse cellular compartments, enforcing the model of a phagosome which interacts extensively with many cellular compartments throughout its progression. Further, our parallel preparation of phagosome samples from human and mouse macrophages enables a species comparison which suggests an increased abundance of antigen preservation and presentation in the human macrophage phagosome at a late stage of maturation. We validate a subset of these observations with orthogonal techniques including organellar cytometry and microscopy. More broadly, this study provides a new methodology which will enable more diverse and unbiased analyses of the phagosome proteome.

## Results

### Analysis of existing phagosome proteomic data yields no high-abundance and phagosome-specific candidate proteins for use as a proximity labeling tether

Proximity labeling has defined the protein composition of various organelles which are difficult to isolate by cell fractionation methods including the intermembrane mitochondrial space, lipid droplets, and the synaptic cleft^49,51,54^. In these studies, compartment-specific proximity labeling was enabled by fusing a proximity labeling enzyme to a high abundance protein “tether” which localizes specifically to that organelle. For example, APEX2 was fused to Perilipin-2, a known lipid droplet protein, for mapping of the lipid droplet proteome by proximity labeling^49^. However, the phagosome is unique in that it is a transient organelle formed first by the plasma membrane then altered throughout its maturation by a series of membrane fission and fusion events with other organelles including endosomes and lysosomes^1^.

We hypothesized that the high overlap between the phagosome and other organelles would result in no endogenous proteins with the high abundance and exclusivity to the phagosome needed to serve as a phagosomal tether (Fig. 1A). To test this hypothesis, we analyzed a previously published phagosome proteomic study by Dill and colleagues which characterized the protein composition of phagosomes isolated from beads conjugated with a variety of ligands at multiple timepoints throughout phagosome maturation^32^. As such, this dataset describes the proteins present at the throughout phagosome maturation across beads conjugated with different ligands to their surface. We searched for an ideal proximity labeling tether for the phagosome meeting three criteria. (1) The protein should localize primarily to the phagosome regardless of the identity of the phagosomal cargo. (2) The protein should persist at the phagosome throughout phagosome maturation. (3) The protein should be transmembrane, as even intact phagosomes can release lumenal content into the cytosol^56^. Of the 1891 proteins quantified by Dill and colleagues, we considered the top 10% most abundant phagosome proteins, and of these 190 proteins, only 12 were transmembrane proteins and maintained at the phagosome throughout phagosome maturation (from 30-minute early phagosomes to 180-minute late phagosomes). 10 of the 12 transmembrane proteins were annotated in the Uniprot database as residing in at least one subcellular compartments other than the phagosome (Fig. 1B). Two proteins, myeloid associated differentiation marker (MYADM) and intercellular adhesion molecule 1 (ICAM1), were the only potentially phagosome-specific nominees, but a literature search revealed that both of these proteins reside abundantly at the plasma membrane^57,58^ and thus would not enable phagosome-exclusive proximity labeling. A recent study utilizing immunoprecipitation for rapid isolation of phagosomes for proteomic and metabolic analyses faced the same challenge and used CD68 as their phagosome tether^59^. However, the authors noted that because CD68 is present throughout endosomes and lysosomes in addition to phagosomes, this strategy cannot definitively distinguish between phagosomes and other endosomes or lysosomes^59^. Based on these observations, we concluded that no known endogenous protein meets the criteria to enable phagosome-specific proximity labeling.

**Figure 1:**
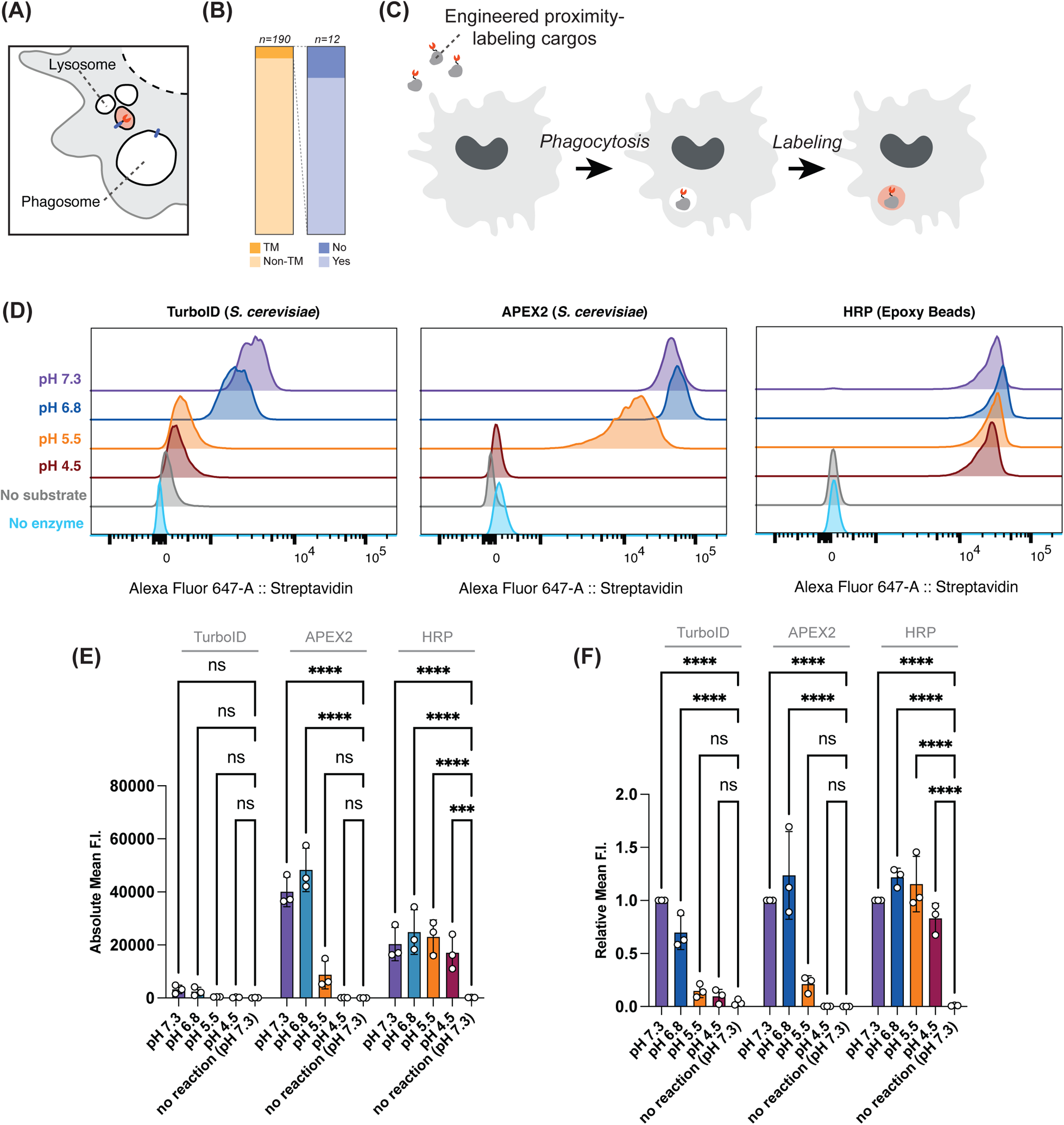
HRP-beads enable proximity labeling across the phagosomal pH range. A Schematic of proximity labeling for membrane bound organelles. A transmembrane protein of a specific cellular compartment is often used as the tether for delivery of a proximity labeling enzyme to a specific compartment. B Analysis of Dill et al. phagosome proteome: of the top 10% most abundant phagosome proteins, 12 proteins were transmembrane (TM) and present at a constant level from early to late phagosome stages. 10 of these 12 proteins were annotated as residing in multiple subcellular locations in the Uniprot database. C Experimental schematic for phagosome lumen proximity labeling mediated by PhagoID. D Representative flow cytometry distributions of TurboID- and APEX2-expressing *S. cerevisiae*, or HRP-conjugated beads reacted with their appropriate labeling substrate across a range of pH, and self-biotinylation was measured by flow cytometry with fluorescent streptavidin staining. E, F Quantification and statistical analysis of (D) across multiple replicates by (E) absolute fluorescence intensity (F. I.) or (F) relative fluorescence intensity (F. I.). (n=3 independent replicates) Data Information: In (E) and (F), data are presented as mean +/− SD. Reported statistics are from a two-way ANOVA with post-hoc comparison of means and Dunnett’s multiple comparisons test; ns indicates adjusted p-value >= 0.05, *** indicates adjusted p-value <0.001, **** indicates adjusted p-value < 0.0001.

### Generating yeast-based and bead-based proximity labeling phagosomal cargoes

In the absence of a high-abundance and phagosome-specific protein, we next hypothesized that phagosomal cargo could be directly imbued with proximity labeling capacity. With this strategy, a proximity labeling enzyme is expressed or otherwise conjugated onto the surface of the phagosomal cargo, and thus the proximity labeling enzyme becomes localized specifically within the phagosome lumen upon phagocytosis of the cargo (Fig. 1C).

There are two modes of proximity labeling commonly used for proteomic analysis: one uses promiscuous biotin ligases (such as TurboID^60^) and the other utilizes peroxidases (APEX2^49–52,61^ and HRP^54,62,63^). In each of these proximity labeling modes, a reactive intermediate (biotin-5′-AMP or phenoxyl free radical for TurboID or APEX2/HRP, respectively) is generated which covalently labels nearby amino acids with a molecular handle for enrichment by affinity purification^64^. We generated three prototypical phagosomal cargos which could catalyze proximity labeling in the phagosome. As APEX2 and TurboID were originally evolved on the surface of *S. cerevisiae*, we generated *S. cerevisiae* capable of expressing either APEX2 or TurboID on their surface^60,65^. APEX2 was evolved in part to overcome a major limitation of HRP which is its inactivity under the reducing conditions of the mammalian cytosol^66,67^. However, because HRP is active in the endosomal and lysosomal compartments and is available as a recombinant protein with high purity, we also generated a bead-based cargo of 3 μm beads with HRP chemically conjugated on their surface^66,68^.

### HRP-bead-based proximity labeling retains its function across the range of phagosomal pH

We hypothesized that the phagosome, which can acidify to a pH of 5 or lower^1,29,30,69–71^, could disrupt proximity labeling reactions. The reported optimal pH of HRP activity ranges from 4.8 to 7^72,73^, and the pH sensitivity of APEX2- and TurboID-catalyzed reactions has never been formally tested. Further, neither APEX2 nor TurboID has demonstrated activity on the lumenal face of the lysosome. As the proximity labeling enzymes and their reaction mechanisms are distinct^60,61,64^, we asked if certain proximity labeling reactions are more resistant to the low pH setting that can be encountered in the phagosome. We tested each of the phagosomal cargos’ ability to label their own surface (self-labeling) across the range of pH that could be encountered within the phagosome^1,29,30,69–71^ (Fig. 1D, 1E, and 1F). For a negative control, we also generated phagosomal cargos with inert proteins on their surface (*S. cerevisiae* surface expressing a non-specific scFv or beads conjugated with BSA). We found that at all tested pH, APEX2-expressing yeast yielded higher levels of self-labeling than TurboID-expressing yeast (Fig. 1D and 1E). When the levels of self-labeling were normalized to the degree of labeling occurring at pH 7.3 (in neat media), APEX2- and TurboID-expressing yeast had significantly reduced labeling at pH 5.5 and lower (Fig. 1D and 1F). It is impossible to directly compare the degree of self-labeling between yeast and beads because of the different particle size and protein content on their surfaces, but we found that the HRP-beads were able to conduct robust labeling across the full range of pH from 7.3 to 4.5 (Fig. 1D, 1E, and 1F). We concluded that HRP-beads are a viable proximity labeling cargo across the pH range that can be encountered within the phagosome at any point throughout its maturation.

### Optimization and development of PhagoID for proteomic analysis of phagosomes with temporal resolution

After confirming that HRP-beads were functional across the phagosomal pH range, we tested that proximity labeling with HRP-conjugated beads could occur in a cellular context. HRP-conjugated beads were added to murine immortalized bone marrow derived macrophages (iBMDMs)^74^ in the presence of the phenolic labeling substrate biotin-phenol (BP) for 45 minutes before adding H_2_O_2_ to catalyze the labeling reaction. The activity of the proximity labeling reaction was measured by probing for protein biotinylation by confocal microscopy (Fig. 2A). We observed biotinylation occurring at a high level only in the presence of BP and H_2_O_2_. Furthermore, this biotinylation was localized specifically to the beads and detectable on fully phagocytosed beads (indicated by white arrows in Fig. 2A; zoomed in field-of-view in Fig. 2B).

**Figure 2:**
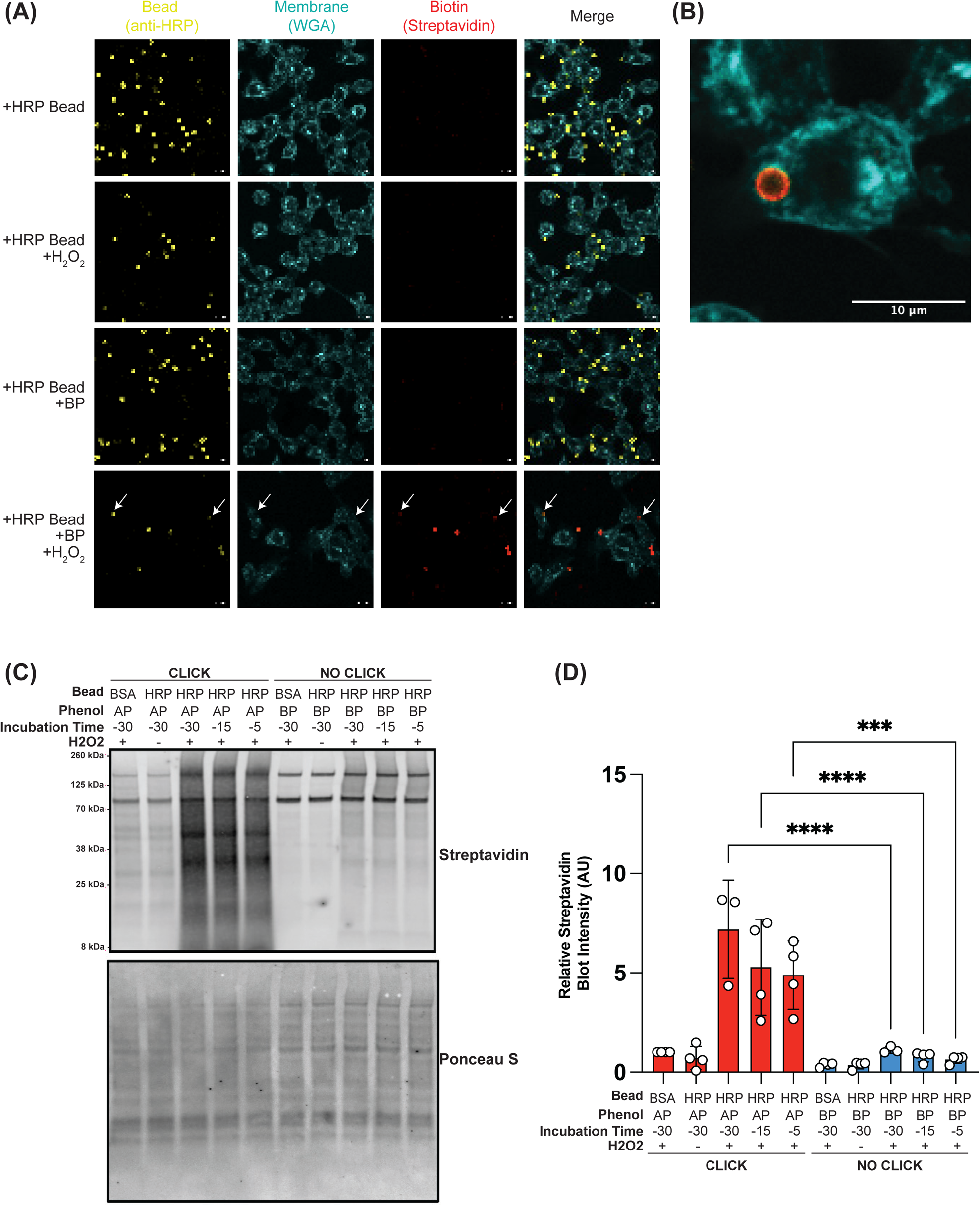
Alkyne-phenol yields more abundant phagosomal labeling than biotin-phenol with an optimized protocol. A iBMDMs phagocytosed HRP-conjugated beads in the presence of 500 μM biotin phenol (BP) for 45 minutes before adding H_2_O_2_ to catalyze proximity labeling and stained with fluorescent streptavidin to visualize abundance and localization of biotinylation in the phagosome by confocal microscopy (white arrows indicate fully internalized beads in the 4th row). Scale bars are 10 μm. B Zoomed in field-of-view of an iBMDM with internalized HRP-bead+BP+H_2_O_2_. Scale bar is 10 μm. C iBMDMs phagocytosed HRP- or BSA-conjugated beads for 30 minutes at 37° C, followed by washing away extracellular beads and another 30 minutes of incubation at 37° C for phagosome progression. The indicated phenolic labeling substrate (500 μM alkyne phenol (AP) or BP) was added post-wash either 30, 15, or 5 minutes prior to labeling. Labeling was catalyzed with H_2_O_2_, click reactions were completed where indicated, and lysates were analyzed by western blot to assess ability of substrate to access the phagosome (representative blot from 3 independent replicates). D Quantification of (C) across independent replicates. Streptavidin band intensities normalized to the Ponceau S intensity indicating total protein abundance (n=3 or 4). Data Information: Reported statistics are from a one-way ANOVA with post-hoc comparison of means and Dunnett’s multiple comparisons test; ns indicates adjusted p-value >= 0.05, *** indicates adjusted p-value <0.001, **** indicates adjusted p-value < 0.0001.

Our microscopy experiments indicated that HRP-conjugated beads could perform proximity labeling in a cellular context. However, based on the resolution of microscopy, we were not able to distinguish between self-biotinylation of the HRP-beads or trans-biotinylation of macrophage proteins. At the same time, we hypothesized that the poor membrane permeability of BP would hinder its localization to the phagosome, reducing the degree of phagosomal proximity labeling^61,75^. We considered an alternative proximity labeling substrate, alkyne-phenol (AP), which has demonstrated more abundant proximity labeling compared to BP in the cytosol of both yeast and mammalian cells, presumably due to its increased membrane permeability^62,75,76^.

To directly observe the trans-labeling of macrophage proteins and test if the improved membrane permeability of AP would increase the abundance of proximity labeling, we used quantitative western blotting. Briefly, HRP-beads (or BSA-beads as a negative control) were added to iBMDMs for an initial period of phagocytosis. Non-internalized beads were washed away and cells were returned to 37° C for phagosome maturation. Either AP or BP was added to cells for 30, 15, or 5 minutes prior to the addition of H_2_O_2_ to catalyze proximity labeling. The labeling reaction was then quenched, cells were lysed, and lysates were prepared to assess the degree of labeling by western blot. AP-based labeling does not result in the direct biotinylation of amino acids, but instead labels with an alkyne modification. Following AP labeling, a copper-based click reaction with biotin-azide conjugates a biotin onto each alkyne modification and enables a direct comparison of the degree of labeling between BP or AP^62,75^ (Fig. 2C and 2D).

By western blot, we detected increased total protein biotinylation in HRP-bead conditions with AP compared to BP regardless of the length of substrate incubation (Fig. 2C and 2D). HRP-beads without H_2_O_2_ and BSA beads with H_2_O_2_ showed minimal protein biotinylation across all conditions. In all lanes we saw the expected bands from endogenously biotinylated macrophage proteins (pyruvate carboxylases at∼130 kDa and CoA carboxylases at ∼70 kDa)^61,77^. To confirm that proximity labeling with AP was not simply more efficient than proximity labeling with BP, we performed proximity labeling with HRP-beads in an axenic setting with either AP or BP. The same degree of bead self-labeling was achieved regardless of the proximity labeling substrate (Supplemental Fig. 1).

Together, these data indicate that using proximity labeling substrates with improved membrane permeability measurably increases proximity labeling at the phagosome. This effect is likely amplified in the PhagoID setting because the labeling substrate must traverse two membranes to reach the phagosome lumen. Further, the membrane permeability of AP makes PhagoID compatible with experimental studies which synchronize phagocytosis by washing away non-internalized beads after an initial period of phagocytosis before adding AP during the secondary period of phagosome maturation^32^.

### Enzymatic digestion of glycoproteins improves quantification of known phagosomal proteins

Our data thus far demonstrated that HRP-conjugated beads were capable of labeling macrophage proteins by proximity labeling in the phagosome. We next sought to utilize our new proximity labeling methodology in a mass spectrometry application to define the protein composition of the phagosome lumen. We hypothesized that many of the proteins labeled by PhagoID would be heavily glycosylated in order to resist the acidic and degradative environment of the phagosome lumen^78^. Glycosylation poses a problem for mass spectrometry, as glycan modifications are variable and complicate database searches of mass spectrometry data^79^. We tested if enzymatic removal of N-linked glycans would augment detection of proteins labeled by PhagoID by eliminating the need to search for the many possible glycan structures at each glycosylation site. We performed the optimized phagosomal proximity labeling in iBMDMs as described (Materials and Methods), then lysates were subjected to a biotin pulldown. Next, each sample was split for one half to be subjected to digestion with peptide-N-glycosidase F (PNGaseF) while the other half remained untreated before samples were prepared for TMT-based quantitative mass spectrometry (Materials and Methods).

Of the 2145 proteins quantified in this experiment, 16 proteins were significantly increased in their quantification upon PNGaseF digestion (Fig. 3A). Of these 16 proteins, 13 were previously confirmed to reside at the phagosome including lysosome-associated membrane glycoprotein-1 (LAMP1), CD63, Procathepsin L (CTSL), and Transmembrane Glycoprotein NMB (GPNMB)^32,43–45^. No other proteins were differentially quantified following PNGaseF digestion (Fig. 3A). As PNGaseF cleaves N-linked glycans from glycosylated asparagines and deamidates the asparagine in the process, we verified that the differential quantitation of these proteins was due specifically to the detection of peptides containing deamidated asparagines which were more abundant in the samples which received PNGaseF digestion (Supplemental Fig. 2). These results demonstrate that PNGaseF digestion enables more reliable detection of glycosylated proteins, and that being able to reliably detect glycosylated proteins increases the accuracy of phagosome proteomics due to the heavily glycosylated proteins in the phagosome lumen.

**Figure 3:**
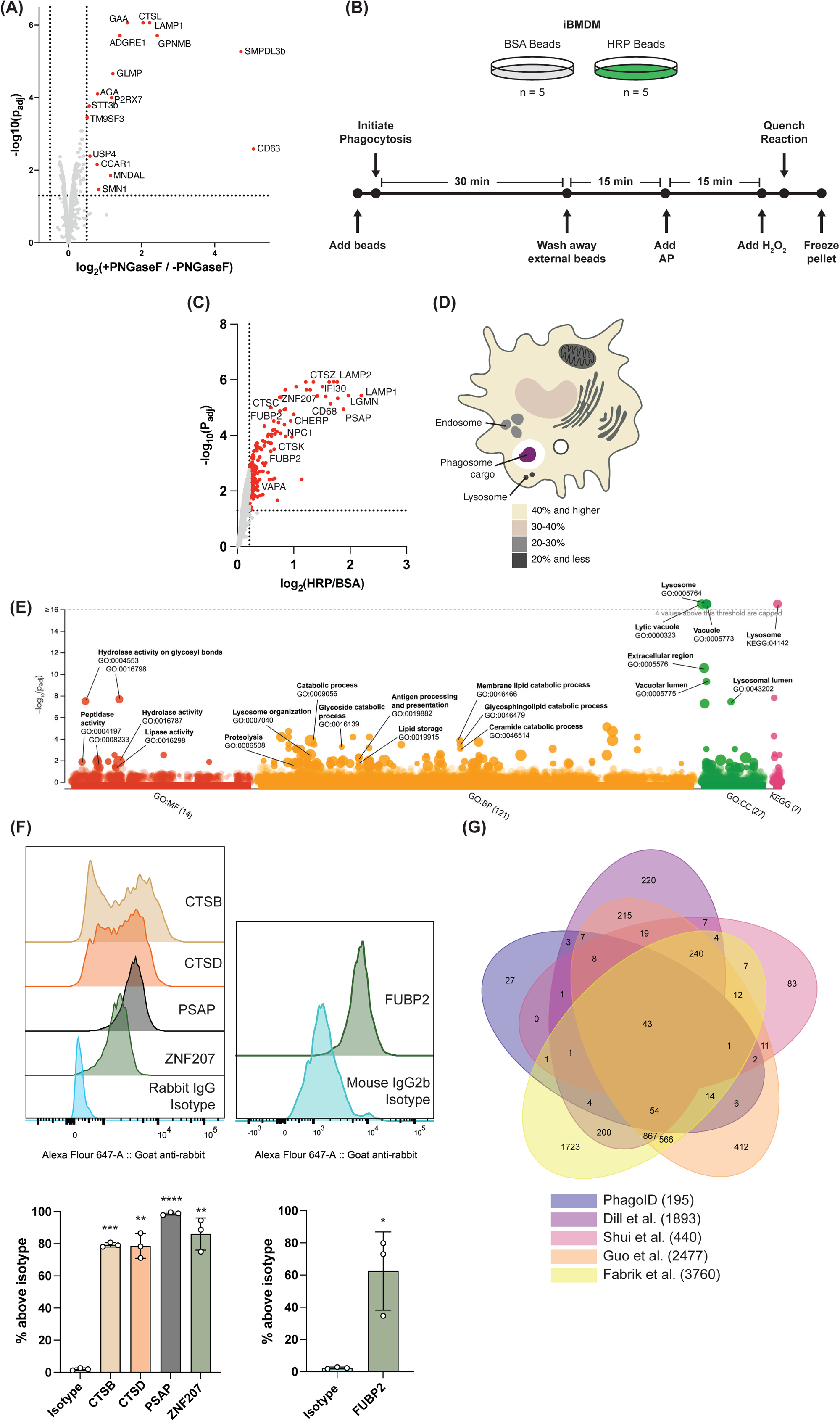
PhagoID enables proteomic analysis of iBMDM phagosomes and highlights nuclear-annotated proteins ZNF207 and FUBP2. A Volcano plot of differentially quantified proteins upon PNGaseF digestion analyzed with the quantitative proteomics workflow (adjusted p < 0.05 and log_2_FC > 0.5). (n=3 independent lysate preparations) B Experimental schematic for TMT-based PhagoID proteomics experiment: 5 replicates were prepared where iBMDMs phagocytosed BSA beads as a negative control, and 5 replicates were prepared where iBMDMs phagocytosed HRP-beads for PhagoID. C Volcano plot highlighting 195 proteins detected at the iBMDM phagosome lumen following statistical ratiometric analysis and thresholding (adjusted p < 0.05 and log_2_FC > 0.21). (n=5 independent lysate preparations) D PhagoID-detected proteins were annotated by their subcellular location according to the Uniprot database and labeled with their percent contribution to the phagosome. E PhagoID-detected proteins were subjected to Gene Ontology (GO) and KEGG enrichment analysis via g:Profiler. All tested terms and p-values are reported in Supplemental Table 2. F Single-phagosome resolution of phagosome colocalization for a panel of PhagoID-detected proteins by PhagoFACS. Representative flow cytometry distributions (upper) and quantification of phagosomes stained above isotype control across multiple replicates (lower). (n=3) G Comparison of significant proteins from PhagoID with previous cell-fractionation-based phagosome proteomic datasets (related to Supplementary Table 3). Data Information: In (A) and (C), p-values were calculated using a moderated T-test and adjusted with the Benjamini-Hochberg protocol. In (E), p-values were calculated using a cumulative hypergeometric test and adjusted with the Benjamini-Hochberg protocol. In (F), data are presented as the mean +/− SD. Where more than 1 antibody was compared to an isotype control, a one-way ANOVA with post-hoc comparison of means and Dunnett’s multiple comparisons test was used. When only 1 antibody was compared to an isotype control, a paired T-test was used. ns indicates adjusted p-value >= 0.05, *** indicates adjusted p-value <0.001, **** indicates adjusted p-value < 0.0001.

### Ratiometric analysis of PhagoID yields a high-confidence proteome of 195 proteins enriched at the iBMDM phagosome lumen

Having established an optimized, robust phagosome proximity labeling workflow compatible with TMT-based quantitative mass spectrometry, we next used this technique to define the composition of the phagosome lumen of murine macrophages. We used our optimized phagosome proximity labeling protocol to determine the protein composition of a mature (60-minute) phagosome (Fig. 3B, Materials and Methods). Our multiplexed mass spectrometry analysis of the murine macrophage phagosome lumen quantified 2536 mouse proteins with 2 or more unique peptides, and there was a high degree of correlation across all five replicates (Supplemental Fig. 3).

We leveraged the quantitative nature of our dataset to apply established analytical workflows to resolve a high-confidence phagosome lumen proteome^50–52,61,62,80^ (Materials and Methods). Although cell fractionation can yield fractions highly enriched for phagosomes, these preparations are never perfectly pure, and the disambiguation of contaminants has complicated previous studies^39,46,81–84^. Most phagosome proteome studies report all of the proteins detected within the phagosome fraction. Our experimental design, in line with standard quantitative proximity labeling protocols^61,62,80^, relies on the comparison between HRP-bead and BSA-bead samples, enabling a ratiometric analysis to filter out proteins which are not heavily enriched at the phagosome (Fig. 3B). We utilized the TMT-based quantitative abundances of each protein to stratify proteins which were specifically labeled by the proximity labeling reaction catalyzed by HRP-beads in the phagosome versus non-specific proteins which are either 1) endogenously biotinylated, 2) non-specific binders to streptavidin beads, 3) non-specifically labeled by click-based biotinylation, or 4) labeled by endogenous peroxidases which catalyze the proximity labeling reaction at sites distal to the HRP-bead. We calculated a normalized fold-change between each protein’s abundance in the HRP-bead sample compared to its paired BSA-bead sample (Materials and Methods). Sorting this list reveals that the proteins enriched in the BSA-bead samples include expected contaminating proteins such as the endogenously biotinylated mitochondrial proteins propionyl-CoA carboxylase and pyruvate carboxylase, as well as the endogenous peroxidase Prostaglandin G/H synthase 1^61,77^. In contrast, the proteins enriched in the HRP-bead samples include known phagolysosomal membrane proteins (LAMP1, LAMP2, GPNMB) and hydrolases (Legumain/LGMN and Prosaposin/PSAP) (Fig. 3C). Thus the TMT-based quantitative proteomics pipeline enables this ratiometric analysis which is critical for distinguishing contaminating proteins from true hits with high confidence.

We used this ratiometric analysis to set statistical and fold-change thresholds to assign proteins as being truly present at phagosome versus non-specific background (Materials and Methods). We set a fold-change threshold to minimize non-specific hits by comparing the dataset to curated true-positive (TP) and false-positive (FP) lists^51,61,62,80^. The TP list is usually defined by a “gold-standard” reference list generated by previous studies. However, because the protein composition of the phagosome lumen is not clearly defined, utilizing a “gold-standard” reference to benchmark our proximity labeling experiments is not possible. Qin and colleagues previously addressed the absence of a “gold-standard” reference by comparing only to a FP list and setting a threshold resulting in a false discovery rate (FDR) of 0.05^62^. We used an identical approach using the Uniprot-curated murine mitochondrial matrix proteome (with proteins cross-listed as “endomembrane network” or “cytoplasm” removed) as the FP list due to its separation from the phagosome lumen by multiple membranes (Materials and Methods). To apply a statistical threshold, we conducted a moderated T-test to identify proteins which were differentially abundant between the BSA-bead and HRP-bead samples, and considered only proteins with a Benjamini-Hochberg adjusted p-value of less than 0.05. Applying the combined statistical and ratiometric abundance filters resulted in a list of 195 murine proteins associated with the phagosome lumen (Fig. 3C, Supplemental Table 1).

### Contribution of endosomes, lysosomes, endoplasmic reticulum, and nucleus to the iBMDM phagosome lumen

Upon mapping the 195 PhagoID-detected proteins to their Uniprot-annotated subcellular locations and gene ontology (GO) analysis, we observed the contribution of multiple subcellular compartments contributing to the phagosome lumen including endosomes, lysosomes, the endoplasmic reticulum (ER), and nucleus (Fig. 3D and 3E).

Many of the 195 proteins were annotated as residing in the endosomes and lysosomes. This included proteins well-characterized to be involved in a variety of lysosomal processes including homo- and hetero-typic endolysosomal fusion (LAMP1, LAMP2)^85^, lipid catabolism (PSAP and ganglioside GM2 activator/GM2a)^86,87^, and protein catabolism (Cathepsins A, B, C, D, K, L, S, and Z)^44,88,89^. The presence of these proteins confirms that the phagosomes labeled by PhagoID have, in bulk, acquired many proteins characteristic of endosomes and lysosomes by the 60-minute timepoint. In total, 56 of the 195 phagosome lumen proteins (28.7%) were annotated as localizing to either the endosome or lysosome (Fig. 3D). In particular, the PhagoID-resolved proteome was heavily enriched for lysosomal proteins (p=7.6×10^−21^; hypergeometric test with Benjamini-Hochberg correction; Fig. 3E), indicating that a substantial degree of phagosome-lysosome fusion has occurred by 60 minutes after internalization.

PhagoID also captured many proteins which highlight the phagosome’s interaction with the ER which has implications for antigen presentation and calcium signaling^90,91^. Vesicle-associated membrane protein-associated protein A (VAPA) was detected and is known to mediate the formation of contact sites between the ER and endosomes, enabling the transfer of cholesterol to modulate membrane morphology and function^92^. Another ER protein, the calcium homeostasis ER protein (CHERP), was also detected. In the ER, CHERP colocalizes with the inositol 1,4,5-trisphosphate receptor (IP3R) where it facilitates Ca2+ release from the ER into the cytosol. CHERP has not been previously detected at the phagosome and its function at the phagosome remains unknown^93^. 17 of the 195 phagosome lumen proteins (8.7%) were annotated as colocalizing to the ER. This does not indicate a significant enrichment relative to the distribution of ER proteins throughout all 2536 quantified murine proteins (p = 0.72; hypergeometric test with Benjamini-Hochberg correction; Fig. 3E). Still, these proteins which at least partially localize to the ER may significantly contribute to the phagosome function.

Several proteins identified by PhagoID were annotated as nuclear proteins that raise new hypotheses about phagosome identity and function. For example, PhagoID identified zinc finger protein 207 (ZNF207), a microtubule-binding protein which localizes to the spindle region and is involved in the separation of chromosomes during cell division^94^. Another set of surprising proteins were the far upstream element-binding proteins 1 and 2 (FUBP1 and FUBP2). In the cytoplasm, FUBP1 and FUBP2 regulate mRNA stability and translation via binding to AU-rich-elements, and have been shown to associate with and disrupt the translation of enterovirus RNA during infection^95^. Splicing factor 1 is another protein detected by PhagoID which is considered to localize primarily to the nucleus and has no characterized function at the phagosome. Of the 195 phagosome lumen proteins, 64 (32.8%) were annotated as localizing to the nucleus. Although this does not equate to a statistical enrichment of nuclear proteins in the PhagoID proteome compared to all 2536 quantified proteins (p = 1; hypergeometric test with Benjamini-Hochberg correction; Fig. 3E), these nuclear proteins were detected as being highly abundant at the phagosome lumen and may play yet unknown roles in phagosome function.

### PhagoFACS validates PhagoID hits and identifies phagosome heterogeneity of cathepsins

To further validate multiple proteins nominated by PhagoID, we conducted phagosome flow cytometry (PhagoFACS)^96,97^. Importantly, our PhagoFACS experiments were conducted with streptavidin-coated dynabeads rather than HRP-coated dynabeads, providing orthogonal validation that these proteins are present at the phagosome independent of a particular protein on the surface of the phagosomal cargo. The lumenal contents of phagosomes can be lost if the phagosome membrane is damaged during the phagosome isolation process. Thus, we only considered phagosomes which had intact membranes at time of analysis to ensure accurate determination of phagosome lumenal contents, as has been done previously^27^. Briefly, phagosomes were isolated from iBMDMs after 60 minutes of phagocytosis with dounce homogenization followed by magnetic isolation. Intact phagosomes were determined by staining with an anti-streptavidin antibody and gating only on phagosomes which remained unstained, indicating the presence of an intact membrane preventing the antibody from binding to streptavidin on the surface of the bead (gating strategy outlined in Supplemental Fig. 4A).

By PhagoFACS of intact phagosomes, we validated the residence of ZNF207, FUBP2, PSAP, and Cathepsins B (CTSB) and D (CTSD) at the 60-minute phagosome (Fig. 3F). ZNF207, FUBP2, and PSAP displayed highly homogenous phagosomal residence, with nearly all 60-minute phagosomes being labeled above the isotype control with the respective antibodies. The transport of PSAP from the Golgi to the phagosome and its residence there has been previously characterized^98^, but this is the first direct validation of ZNF207 and FUBP2 at the murine phagosome.

In contrast, we observed marked heterogeneity in the phagosome residence of CTSB and CTSD, indicated by the bimodal distribution of CTSB and CTSD abundance on individual phagosomes 60-minutes after internalization (Fig. 3F). Notably, this heterogeneity was consistent across all replicates (Supplemental Fig. 4B). The heterogeneous distribution of CTSB and CTSD throughout endosomal and lysosomal populations has not been characterized, to the best of our knowledge. The observed bimodal distributions of CTSB and CTSD observed at the phagosome at 60-minutes post-internalization may suggest that there is regulated transfer of these cathepsins to the phagosome at this point in phagosome maturation.

### Comparing multiple phagosome proteomic studies yields a consensus core phagosome lumen proteome

In the absence of a gold standard phagosome lumen protein composition, we reasoned that comparing across multiple phagosome proteome mass spectrometry datasets would construct a gold standard phagosome lumen proteome. We compared the proteins identified by PhagoID with four other proteomic analyses of the bead-containing phagosome (Fig. 3G; Supplemental Table 3). Collectively these studies span a variety of murine macrophage cell types, phagosome maturation timepoints, phagosome isolation methods, and sample processing to either capture the entire phagosome^32,44,45^ or enrich for the phagosome membrane^43^ or lumen (PhagoID). Spanning the heterogeneity of these approaches, there is a consensus core phagosome lumen proteome consisting of 43 proteins agreed upon by all five compared proteomic studies (Fig. 3G; Supplemental Table 3). GO and KEGG enrichment analysis of the core phagosome lumen proteome highlights an enrichment for hydrolase activity, lipid metabolism, and antigen presentation (Supplemental Fig. 5).

Of the 43 core phagosome proteins which share a consensus among all five datasets, 11 proteins are hydrolases responsible for the catabolism of proteins, glycans, lipids, and nucleic acids such as cathepsins, beta-hexosaminidase, beta-galactosidase, saposins (collectively detected by mass spectrometry as PSAP), and the PLD3 and PLD4 exonucleases. Two proteins in the core phagosome proteome, Ferritin and NPC Intracellular Cholesterol Transporter 1 (NPC1), highlight the phagosome’s role in processing, storing, and transporting nutrients. NPC1 functions in tandem with NPC2 to transport cholesterol out of the phagosomal compartment^9,99^. Ferritin is responsible for storing iron and functions not only to recycle nutrients from phagocytosed cargo, but also starves any pathogens in the phagosome by sequestering iron^100^. Clearly, the core phagosome proteome highlights the degradative nature of the phagosome, and its essential function in catabolizing phagocytosed targets to simple molecules for their recycling.

Surprisingly, there remain several proteins identified among the core phagosome proteome whose phagosomal function remains unclear. For example, Splicing factor 1 is involved in the assembly of the spliceosome in the nucleus to remove introns from pre-mRNA before mRNA export into the cytosol^101^. Its presence in the core phagosome proteome is surprising, as its function outside of the nucleus is not characterized. Additionally, SLC25A5 is a transmembrane transporter of ADP and ATP into and out of the mitochondrial matrix, but its function at non-mitochondrial membranes is not defined^71,102^. Acid sphingomyelinase-like phosphodiesterase 3b (SMPDL3b) is another member of the core phagosome proteome whose function at the phagosome is yet unknown, but has been characterized as a lipid-modifying enzyme which localizes to the plasma membrane and modulates TLR signaling upon stimulation with purified lipopolysaccharide (LPS) or E. coli *in vitro* and *in vivo*^103^.

Of interest were the 240 proteins that shared consensus from the other 4 datasets but were undetected by PhagoID. Included among these 240 proteins were 11 Rab proteins, which are known to scaffold exclusively on the outer membrane of phagosomes to regulate phagosome trafficking^6,12,26,27,104^. The absence of these proteins from the PhagoID-resolved lumenal proteome confirms that PhagoID enables proteomic analysis with topological specificity and enriches for lumenal phagosome proteins. Also among these 240 proteins not captured by PhagoID were many subunits of the vacuolar ATPase which acidifies the phagosome lumen.

Of the 4781 proteins identified at the phagosome across the five considered proteomic datasets, 27 of them were uniquely detected by PhagoID (Fig. 3G). This group includes proteins spanning a wide variety of activity including nucleic acid sensing and degradation (RNAse-T2b and DCP1a), aminopeptidase activity (NPEPL1 and XPNPEP1), lipid remodeling (GBA1 and HMGCS1), ubiquitination and autophagy (UBE2E1 and ATG2a), as well as several poorly characterized proteins with unclear functions (e.g. MAK16 Homolog, Small acidic protein, HEATR6). Although many factors may contribute to other studies not detecting these proteins, it is possible that these proteins are lowly abundant and require the specific enrichment labeling of lumenal proteins for their detection. The differential abundances of these novel proteins in the BSA vs. HRP conditions are shown in Supplemental Figure 6.

### PhagoID resolves the primary human macrophage phagosome lumen with limited cellular input

As proper phagosome function is critical for maintaining human health, and discrepancies in the biochemical environment between human and murine phagosomes have been previously described^105^, we sought to measure the phagosome lumen of a primary human macrophage. The large number of cells required for fractionation-based phagosome proteomics has previously limited the analysis of primary human phagosomes^46,47^. PhagoID directly addresses this limitation, as we completed our iBMDM PhagoID experiment utilizing ∼50e6 macrophages per sample compared to previous fractionation-based studies which required ∼700e6 macrophages per sample^32^. Additionally, primary human phagocytes are difficult to genetically engineer and have not been subject to previous proximity labeling studies. Thus we reasoned that PhagoID, which requires no genetic engineering and is achievable with reduced cell input, would uniquely enable us to define the phagosome lumen proteome of a primary human phagocyte where ongoing knowledge gaps exist.

To define the composition of the phagosome lumen in primary human monocyte-derived macrophages (hMDMs), we first generated hMDMs from positively-selected CD14+ monocytes from five healthy human donors. For each sample, 25e6 monocytes were differentiated in M-CSF for 6 days as we have previously^106,107^. Following differentiation, we pursued an identical experimental approach as with iBMDMs (Materials and Methods). Given that these are primary human cells derived from different donors, we did observe greater variation across these samples than our murine macrophage experiments (Supplemental Fig. 7). Our quantitative multiplexed mass spectrometry analysis of the primary hMDM phagosome lumen identified 1016 human proteins with 2 or more unique peptides across all BSA- and HRP-bead conditions. We hypothesized that the curated core phagosome lumen to be conserved at the hMDM phagosome lumen, so we utilized the human orthologs of the 43 core proteins (Fig. 3G) as a TP list. We again used the Uniprot-curated human mitochondrion matrix proteome (with proteins cross-listed as “endomembrane network” or “cytoplasm” removed) as an FP list due to its separation from the phagosome by multiple membranes. We observed that the core phagosome lumen proteome was enriched relative to the proteins of the mitochondrial matrix (Fig. 4A and Supplemental Fig. 7). We then selected the fold-change cutoff which maximized the difference between the TP rate and the FP rate (Supplemental Fig. 7, Materials and Methods). We again applied a statistical threshold by conducting a moderated T-test between the HRP-bead and BSA-bead conditions and considering only proteins with a Benjamini-Hochberg adjusted p-value of less than 0.05. Applying the combined statistical and ratiometric abundance filters resulted in a list of 123 human proteins associated with the phagosome lumen (Fig. 4B; Supplemental Table 4).

**Figure 4:**
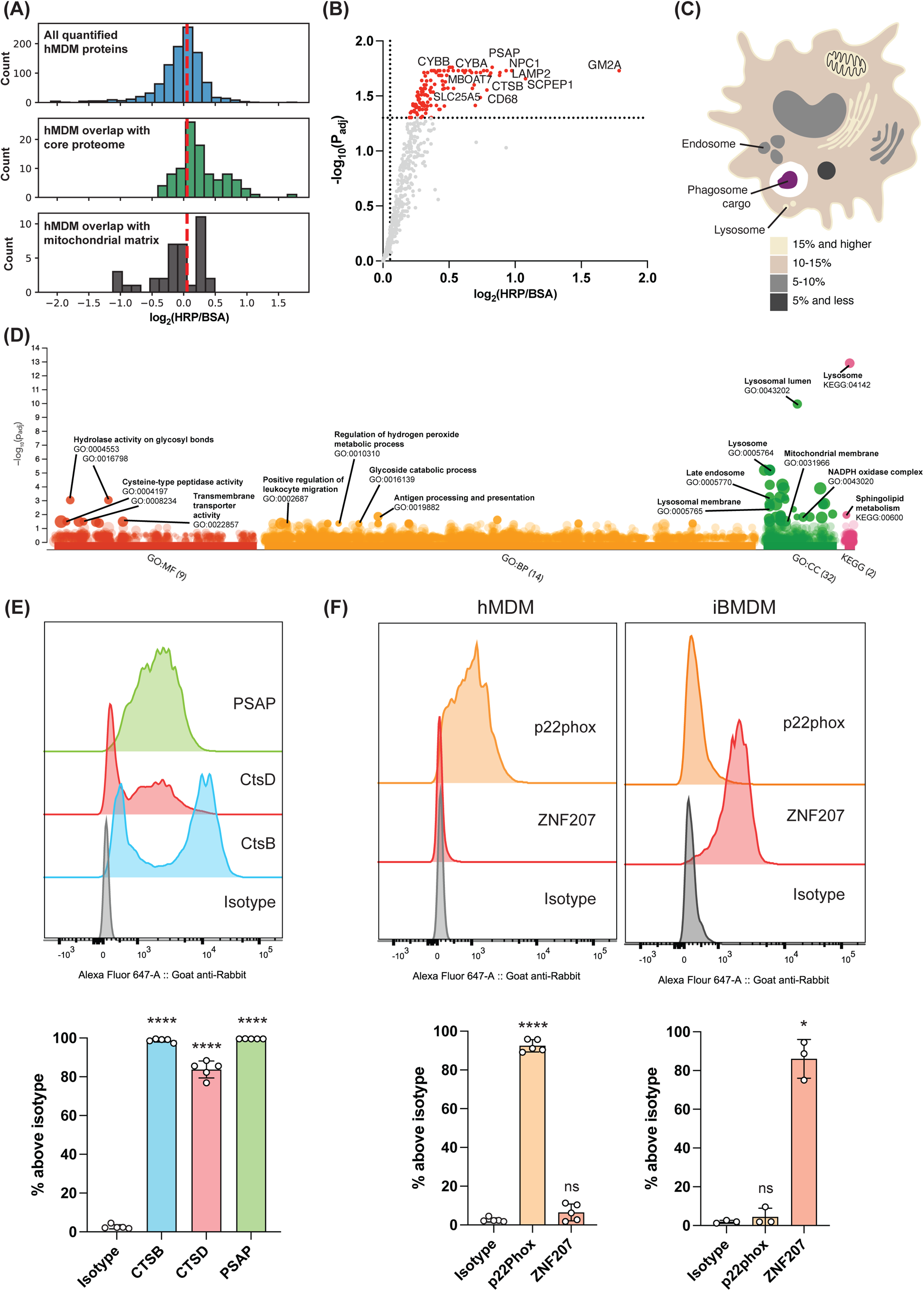
PhagoID enables proteomic analysis of primary human monocyte-derived macrophages and identifies increased abundance of the NADPH oxidase complex. A Histogram depicting hMDM phagosome lumen quantified log2-foldchange distribution of all proteins (top), proteins among the core phagosome lumen proteome (middle), and proteins among the FP list mitochondrial matrix (bottom). The applied fold-change threshold is indicated with a dashed line. B Volcano plot highlighting 123 proteins detected at the hMDM phagosome following statistical ratiometric analysis and thresholding (adjusted p < 0.05 and log_2_FC > 0.0538). (n=5 human donors) C PhagoID-detected proteins were annotated by their subcellular location according to the Uniprot database and labeled with their percent contribution to the phagosome. D PhagoID-detected proteins were subjected to Gene Ontology (GO) and KEGG enrichment analysis via g:Profiler. All tested terms and p-values are reported in Supplemental Table 2. E Single-phagosome resolution of phagosome colocalization for a panel of PhagoID-detected proteins by PhagoFACS. Representative flow cytometry distributions (upper) and quantification of phagosomes stained above isotype control across multiple replicates (lower). (n=5 human donors) F PhagoFACS validation of proteins nominated by PhagoID for differential abundance in hMDM and iBMDM. Representative flow cytometry distributions (upper) and quantification of phagosomes stained above isotype control across multiple replicates (lower). (n=5 human donors for hMDM and n=3 for iBMDM). **Same data replotted for iBMDM ZNF207 staining as from Fig. 3F for side-by-side comparison with hMDM**. Reported statistics from where more than 2 categories are compared are from a one-way ANOVA with Dunnett’s multiple comparisons test. Where 2 categories are compared, a Paired T-test was used. ns indicates adjusted p-value >= 0.05, *** indicates adjusted p-value <0.001, **** indicates adjusted p-value < 0.0001.

### Contributions of endosomes, lysosomes, and mitochondria to the hMDM phagosome lumen

By comparing the hMDM PhagoID proteins to the Uniprot database and GO analysis, we observed many proteins at the hMDM phagosome lumen annotated as colocalizing to multiple different cellular compartments including the endosomes, lysosomes, and mitochondria (Fig. 4C and 4D). In total, 37 of the 123 hMDM phagosome lumen proteins (30.1%) were annotated as localizing to either the endosome or lysosome (Fig. 4C). Many of the same characteristic endosomal and lysosomal proteins were detected as were in iBMDMs, including lysosomal transmembrane proteins (LAMP2 and SCARB2), lipid catabolism (PSAP and GM2a), protein catabolism (Cathepsins A, B, D, H, L, S, and Z), and nucleic acid catabolism (PLD3 exonuclease). The hMDM PhagoID-resolved proteome was heavily enriched for lysosomal proteins (p = 6.6×10^−6^; hypergeometric test with Benjamini-Hochberg correction; Fig. 4D) indicating that substantial phagosome-lysosome fusion has occurred by 60 minutes after internalization. Clearly, the 60-minute hMDM phagosome acquired many degradative functions to facilitate the breakdown of its phagosomal cargo, as is corroborated by the enrichment of several degradative molecular functions and biological processes by GO analysis (Fig. 4D). We validated the presence of PSAP, CTSB, and CTSD by PhagoFACS of intact hMDM phagosomes. PhagoFACS analysis of hMDM phagosomes revealed similar patterns of heterogeneity in the acquisitions of CTSB and CTSD to that seen in iBMDM phagosomes, and the observed heterogeneity was consistent across all human donors (Fig. 4E, Supplemental Fig. 4C).

24 of the 123 hMDM phagosome lumen proteins (19.5%) were annotated as residing in the mitochondria (Fig. 4C). The mitochondrial superoxide dismutase (SOD2) was one protein detected by hMDM PhagoID. SOD2 is primarily responsible for destroying H_2_O_2_ produced in the mitochondrial matrix, but was recently shown to be transported from the mitochondria to bacteria-containing phagosomes by mitochondria-derived vesicles^108^. SLC25A5, discussed above, was also detected alongside two other mitochondrial inner membrane transporters SLC25A3 and SLC25A11. Similar to SLC25A5, SLC25A3 and SLC25A11 are both proteins whose presence at the phagosome is corroborated by multiple phagosome proteomic studies but whose non-mitochondrial function is not well characterized^32,43–45,71^. In accordance with these findings, the hMDM phagosome lumen proteome was enrichment for the GO terms “mitochondrial membrane” and “mitochondrial envelope” (p = 0.032 and 0.021; hypergeometric test with Benjamini-Hochberg correction; Fig. 4D). However, the general term “mitochondria” was not enriched (p = 0.35; hypergeometric test with Benjamini-Hochberg correction; Fig. 4D).

### Species comparison of phagosome proteomes reveals an increase in NADPH oxidase complex and antigen presentation machinery at the hMDM phagosome lumen

Because the phagosome preparations from iBMDM and hMDM were prepared identically in this study, we reasoned that we could gain insight into distinct phagosomal features between the macrophages derived from two different species. We mapped each detected murine protein to its human ortholog where possible, then compared the overlap of the 123 hMDM phagosome lumen proteins with the 195 iBMDM phagosome lumen proteins. Of these proteins, 39 overlapped between both cell types while 84 were unique to hMDMs and 156 were unique to iBMDMs.

Many of the 39 overlapping proteins are characteristic lysosomal proteins discussed previously, such as LAMP2, LAMP4, GPNMB, cathepsins, NPC1, and PSAP. Coronin-1C, a protein involved in actin dynamics throughout phagosome formation and early phagosomal budding events, was also detected across both species^109^.

Interestingly, both of the lumen-accessible subunits of the membrane-associated NADPH oxidase (p22Phox/CYBA and gp91Phox/CYBB), were detected as significantly abundant in the hMDM phagosome proteome, but not in the iBMDM phagosome proteome. Further, multiple components of antigen processing machinery including both class I and II MHC molecules (HLA-A, HLA-DRB5, HLA-DPB1), Antigen Peptide Transporter 1 (TAP1), and the ER aminopeptidase (ERAP1) were also detected by hMDM PhagoID, but the equivalent murine proteins were not detected in the iBMDM PhagoID experiment. The differential abundance between iBMDM and hMDM for each of these proteins as quantified by mass spectrometry is plotted in Supplemental Figure 8A.

We used antibodies with reported cross-reactivity to both human and murine proteins to validate two proteins, CYBA and ZNF207, which were indicated by PhagoID as being much more abundant in either hMDM or iBMDM phagosomes. We observed that nearly all hMDM phagosomes stained positive for CYBA at the 60-minute timepoint while there was essentially no signal above isotype control present on iBMDM phagosomes (Fig. 4F). Upon staining phagosomes for ZNF207, we observed that ZNF207 was abundant at nearly all iBMDM phagosomes, with no signal above isotype control on hMDM phagosomes (Fig. 4F). Interestingly, Western blotting of whole-cell lysates shows that CYBA, but not CYBB, is less abundant overall in iBMDMs. ZNF207 is present in higher abundance in iBMDMs, but is still readily detectable in hMDMs (Supplemental Fig. 8B and 8C). These findings corroborate our PhagoID experiments, as ZNF207 was detected by PhagoID in iBMDMs but not hMDMs, and CYBA was detected in hMDMs but not iBMDMs.

## Discussion

The phagosome is the organelle which enables professional phagocytes to maintain homeostasis in a variety of contexts: from clearing apoptotic cells to tissue remodeling to killing pathogens and infected cells. The diversity and adaptability of the phagosome is encoded in its protein composition. Here we developed a new technique to simplify the analysis of this organelle that is critical for immunity by establishing a modular proximity labeling strategy for phagosome proteomic analysis.

We enabled proximity labeling exclusively at the phagosome by conjugating HRP onto the surface of a phagosomal cargo and developed an optimized protocol for phagosomal proximity labeling. We utilized this approach to characterize the phagosome lumen proteomes of immortalized murine macrophages and primary human macrophages with TMT-based quantitative mass spectrometry. Importantly, we were able to generate a list of high-confidence proteins enriched at the bead-containing phagosome lumen by filtering out non-specific proteins using the ratiometric analysis enabled by proximity labeling. Due to the specific targeting of the phagosome lumen and ratiometric filtering of the data, the PhagoID phagosome lumen proteome is a smaller list than previous phagosome proteomes^32,43–45^, consisting of high-confidence proteins with specific enrichment to the phagosome lumen. Our PhagoID-resolved proteomes of iBMDMs and primary hMDMs largely agree with previous fractionation-based datasets^32,43–45^, but PhagoID also nominates novel phagosome lumen constituents. Consistent with the highly interactive nature of the phagosome, PhagoID identified many proteins annotated across multiple cellular compartments beyond the “main” contributors such as plasma membrane, endosomes, and lysosomes.

We detected and validated the presence of two previously unvalidated and nuclear-annotated proteins at the murine phagosome: FUBP2 and ZNF207. In the nucleus, FUBP1 and FUBP2 (also known as KHSRP) enforce epigenetic regulation of MYC expression levels ^110^. In the cytoplasm, FUBP1 and FUBP2 regulate mRNA stability and translation via binding to AU-rich-elements. The exact signal which modulates FUBP localization is not known, but FUBP2 has been shown to relocalize to the cytoplasm upon enteroviral infection of human rhabdomyosarcoma cells where it disrupts the translation of enteroviral RNA during infection^95^. The detection of FUBP1 and FUBP2 suggests that FUBP1 and FUBP2 gain access to the phagosome following phagocytosis, raising potential implications for their involvement in surveillance of the phagosome for viral RNAs. ZNF207 is a DNA-binding and microtubule-binding protein with seemingly dual functions in the nucleus and cytoplasm. In the nucleus, ZNF207 regulates the self-renewal capabilities of stem cells^111^. ZNF207 is localized exclusively to the nucleus during interphase, then distributed throughout the cytosol following nuclear envelope breakdown where it localizes to the spindle region throughout the rest of the cell cycle^94^. Given that ZNF207 is involved in regulation of the cell cycle, we considered that the presence of ZNF207 could be an artifact from using an immortalized cell line, however ZNF207 was also detected in a proteomic analysis of magnetic-bead containing phagosomes derived from primary murine tissue resident macrophages^44^. Interestingly, ZNF207 was detected specifically at the iBMDM phagosome and not at the hMDM phagosome. The association of ZNF207 with the phagosome lumen and its functional impact at the phagosome remain an area for further study.

Multiple interesting proteins which typically reside at the mitochondria were detected at the phagosome, highlighting an increasingly studied field of communication between the endosomal compartment and mitochondria^108,112,113^. In particular, SOD2 was one mitochondrial protein detected by both iBMDM and hMDM PhagoID. Intriguingly, SOD2 has been shown to be delivered to the bacteria-containing phagosomes via mitochondrial-derived vesicle fusion, but this was not observed with bead-containing phagosomes^108^. Mitochondria-phagosome juxtaposition has also been demonstrated at bead-containing phagosomes following TLR stimulation to support the phagosomal oxidative burst, but the transfer of proteins to the phagosome in this scenario was not directly observed^112^. Another possible cause of mitochondrial contribution to the phagosome is mitophagy, in which mitochondria are taken up into lysosomes for degradation, followed by phagosome fusion with the mitochondria-containing lysosome^113,114^. In this scenario it would be expected that the mitochondrial proteins detected at the phagosome would largely be the most highly abundant mitochondrial proteins. However, SOD2 is detected at relatively low levels within the mitochondrial proteome^36^. Thus in our study the mechanism of SOD2 delivery to the phagosome remains unclear, as we observed SOD2 at the bead-containing phagosome in the absence of any TLR stimulus.

PhagoID enabled a comparison between the phagosome lumen of iBMDMs and hMDMs. Prior to this, no study has prepared phagosomes derived from the two species for an identical proteomic analysis. PhagoID detected an increased abundance of the two lumen accessible components of the NADPH oxidase complex (p22Phox/CYBA and gp91Phox/CYBB), as well as increased antigen presentation machinery on hMDM phagosomes compared to iBMDM phagosomes. The NADPH oxidase complex is responsible for generating the oxidative burst within the phagosome, which serves to both kill phagocytic cargo with reactive oxygen species and preserve its antigens for presentation by limiting the rate of acidification and proteolysis in the phagosome^115,116^. Interestingly, a previous study also identified the NADPH oxidase subunit CYBB as being differentially transcriptionally regulated between human and murine macrophages following stimulation with LPS^117^. However, in this study human CYBB mRNA decreased in expression following stimulation with LPS whereas murine CYBB mRNA increased following LPS stimulation^117^. Thus, the divergent regulation of NADPH oxidase mRNA and protein between murine and human macrophages remains an area for future study. However, the increased abundance of NADPH oxidase and antigen presentation proteins likely indicate that the hMDM phagosome retains a higher propensity for antigen presentation at this timepoint.

We also compared multiple proteomic analyses and compiled a “core phagosome lumen proteome” corroborated by 5 phagosome proteome datasets. Many of the proteins belonging to this core proteome have well-annotated functions as endosomal membrane proteins or lysosomal catabolic enzymes, but this analysis also revealed several surprising proteins which have been detected by many proteomic studies, but whose function at the phagosome is not yet characterized: Splicing Factor 1, SMPDL3b, and SLC25A5. Based on their annotated functions, we can speculate that these proteins may contribute at the phagosome towards nucleic acid processing, lipid remodeling, and ATP / H+ transport, respectively. As these proteins are highly corroborated at the phagosome, further characterization of the roles that these proteins play at the phagosome and their impact on phagosome function is an exciting area for future studies.

Taken together, these data demonstrate the utility of PhagoID and proximity labeling to quantitatively resolve the proteome of the phagosome lumen with high confidence, granting clear insight into phagosome identity and nominating proteins for further study to gain novel insight into the phagosome function. Further, this study provides an optimized protocol for proximity labeling in the phagosome and serves as a template for future studies to measure and compare the phagosome lumen proteome across a variety of phagosomal cargos and cell states.

Although we chose to conjugate HRP onto the surface of beads to enable a direct comparison between PhagoID and established fractionation-based methods, this approach can be applied to many phagosomal cargoes such as bacteria or apoptotic cells and enables the direct comparison of protein composition through ratiometric analysis. For example, a recent study published by Li and colleagues utilized APEX2 tethered to the surface of multiple microbes to capture microbe-specific phagosome proteomes measured by label-free quantitative mass spectrometry^118^. In their phagosomal proximity labeling protocol, they did not wash away extracellular proximity labeling cargo but instead quenched any extracellular labeling reactions by adding a membrane-impermeant antioxidant epigallocatechin-3-gallate (EGCG) immediately before adding H2O2. We propose that an optimized protocol would be a hybrid of our approaches, utilizing AP instead of BP and applying EGCG to fully prevent any extracellular labeling. With this protocol, washing away extracellular beads is only required in cases where synchronized phagocytosis is desired.

PhagoID also theoretically enables the proteomic analysis of damaged phagosomes which are incompatible with the fractionation-based approach on account of their damaged membranes causing the loss of lumenal contents and dissociation from the phagosomal cargo during homogenization and centrifugation. However, because HRP is active only within the secretory system and inactive under the reducing conditions of the cytosol, conditions with high degrees phagosomal damage which could alter the reducing environment of the phagosome may be better served by APEX2-based PhagoID. Further, the low cell input required for PhagoID enables comparisons of phagosome proteomes derived from cells which are not available in high abundances, such as primary cDC1s or primary human tissue resident macrophages which can only be isolated in relatively small numbers^119,120^. The application of PhagoID to each of these areas: diverse phagosomal cargo, previously inaccessible phagosome conditions, and rare cell types, presents an exciting future in which we can better measure and understand how the phagosome proteome is shaped in a variety of contexts.

### Limitations of this study

In this study, we utilized proximity labeling combined with TMT-based quantitative mass spectrometry for identification and quantification of phagosomal proteins. As are all proximity labeling studies, this study is intrinsically limited by the ability of peroxidase-mediated proximity labeling to only label accessible tyrosine residues^64^. Thus, any protein without tyrosines available to the phagosome lumen may not be detected by PhagoID. Additionally, peroxidase-based proximity labeling (HRP and APEX2) is widely used in cell culture applications, but is not easily adapted to *in vivo* applications due to the requirement for H_2_O_2_. Although we observed lower levels of labeling by TurboID on the surface of *S. cerevisiae*, proximity labeling catalyzed by TurboID-engineered cargo may be sufficient to label proteins in the phagosome, provided sufficient availability of biotin and ATP in the phagosome. However, we observed reduced TurboID activity at lower pH. Thus, any TurboID-engineered cargo should be benchmarked for activity across a pH range to understand if the proximity labeling reaction is occurring in all phagosomes, or only early phagosomes which have not yet fully acidified.

We chose to utilize TMT-based quantitative mass spectrometry as it enables high-confidence and directly comparable quantification of proteins across 10 samples, but it has been shown that TMT-based approaches yield lower proteome coverage than label-free quantitative mass spectrometry^121^. Further, while the ratiometric comparison of PhagoID samples to negative controls filters out many common contaminants, it is possible that some proteins which are lowly abundant, or present at the phagosome but not highly enriched relative to the whole-cell proteome, were filtered out. Our comparisons of proteins enriched in the hMDM phagosome lumen to those enriched in the iBMDM lumen are limited by the fact that these samples were run separately rather than in the same TMT multiplexed analysis. Selection of peptides for acquisition in separate TMT multiplexed analyses is inherently variable. Some orthologous proteins may have been detected in one experiment but not the other for technical rather than biological reasons (for example, sequence differences causing tryptic peptides to have different ionization efficiency).

As we identified a total of 279 proteins at the hMDM and iBMDM phagosomes, we were unable to validate all of these proteins with a secondary method. One major limitation of validation studies is the availability of high-quality antibodies that are suitable for intracellular protein staining for microscopy and flow cytometry. In particular, the reduced availability of antibodies which were reported as being cross-reactive to both human and mouse limited our ability to compare differences between hMDM and iBMDM. Even when an antibody is cross-reactive, there is no guarantee that the antibody has the same affinity for the same protein from the two species if there are any differences in the amino acid sequence between the two species.

## Materials and methods

### Preparation of BSA- and HRP-beads

HRP and BSA were conjugated to epoxy surface-functionalized Dynabeads using the Antibody Coupling Kit (ThermoFisher, 14311D) according to the manufacturer’s protocols, with the following modifications: 150 ug of protein per 1 mg of beads was added to the bead/protein mixture, and conjugation was carried out at 4° C for 72 hours before washing. Each bead was washed in appropriate cell culture medium 3x before use.

### Preparation of APEX2- and TurboID- surface expressing yeast

pCTCON2 template plasmid (a gift from Dane Wittrup) was modified to contain the coding regions for APEX2-V5 and TurboID-V5 to the C-terminus of Aga2p. Yeast were transformed using the Frozen EZ-yeast Transformation II Kit (Zymo Research, T2001) according to the manufacturer’s protocols. Surface display was induced by inoculating yeast cultures into SDCAA with shaking at 30° C until an OD of 0.5-1.0 was reached. Surface display induced with SD/GCAA containing a mixture of 10% dextrose and 90% glucose with shaking at 20° C for 18-48 hours. TurboID yeast were grown in biotin-depleted variations of these media, prepared with biotin-depleted yeast nitrogen base (Sunrise Science, 1523-100) to reduce background labeling.

### pH titration and assessment of self-labeling

After overnight induction, yeast OD was measured and 2e6 yeast were prepared for each sample (assuming 30e6 yeast/mL per OD unit). Similarly, beads were counted and 2e6 beads were prepared for each sample. DMEM with 10% FBS (D10) was equilibrated to atmospheric CO2 at 37° C, then acidified to the desired pH with concentrated HCl. Yeast or beads were washed 3x with the appropriate pH media, then resuspended in 100 uL of the appropriate pH media in the presence or absence of labeling substrate (250 μM BP (Iris Biotech, LS-3500) for APEX2/HRP or 250 μM biotin + 100 μM ATP for TurboID) as indicated. Yeast or beads were incubated in acidified D10 at 37° C with shaking for 10 minutes to simulate cell culture conditions. For TurboID conditions, yeast were washed after the 10 minute labeling window with 3x with ice-cold DPBS. For APEX2-yeast and HRP-beads, H_2_O_2_ was added to a final concentration of 1 mM with gentle agitation for 60 seconds, then the reaction was quenched by adding an equal volume of ice-cold 2X quenching buffer (DPBS containing 10 mM trolox (Cayman Chemical, 10011659), 20 mM sodium ascorbate (Sigma, A4034), 10 mM sodium azide (Sigma, S2002)). APEX2-yeast and HRP-beads were then washed 3x with ice-cold quenching buffer (DPBS containing 5 mM trolox, 10 mM sodium ascorbate, 10 mM sodium azide). Prior to staining, all samples were washed 2x with ice-cold 1% BSA, and staining was completed in 1% BSA.

For yeast, surface expression levels were measured by staining with a rabbit anti-V5 antibody for 30 minutes on ice (1:1000, CST, #13202), followed by three washes and subsequent staining with an AlexaFluor 488-conjugated goat anti-rabbit antibody on ice for 30 minutes (1:1000 dilution, ThermoFisher, A-11008). Self-labeling of yeast and beads was measured by staining with AlexaFluor 647-conjugated streptavidin (4 ug/mL, ThermoFisher, S21374) on ice for 30 minutes. Samples were washed 3x after staining and resuspended in 1% BSA for analysis on an a for analysis on an LSRFortessa flow cytometer (BD Life Sciences).

All flow cytometry data was analyzed using FlowJo™ v10.8 Software (BD Life Sciences). Yeast were gated to only include populations with equal levels of surface-display expression for analysis. To calculate relative fluorescence where indicated, all values for a replicate were internally normalized by dividing the mean absolute fluorescence intensity for each sample by the mean absolute fluorescence intensity of the labeling reaction conducted at pH 7.3 for that replicate.

### Confocal microscopy of proximity labeling with HRP-beads

For confocal microscopy, 75k iBMDMs were seeded per well into a 12-well coverslip (Ibidi, 81201). The following day, 500k of the appropriate beads (HRP- or BSA-beads) were prepared with 3x washes of D10 then added per well in the presence or absence of 500 μM BP. After 45 minutes of phagocytosis, H2O2 was added to a final concentration of 1 mM with gentle agitation for 60 seconds, then the reaction was quenched by adding an equal volume of ice-cold 2x quenching buffer. Cells were washed 3x with 1x quenching buffer, then fixed for 10 minutes with methanol at −20° C. Cells were then washed 3x with DPBS and further permeabilized and blocked by incubation with blocking buffer (DPBS +1% BSA +0.3% Triton X-100) for 1 hour at room temperature. Cells were stained with the primary anti-HRP mouse antibody diluted 1:1000 in blocking buffer overnight at 4C. The following day, cells were washed 3x with DPBS (3 minutes per wash), then stained with streptavidin-AF647 (2 ug/mL), DAPI (1 ug/mL), WGA-AF488 (5 ug/mL, ThermoFisher, W11261) and Alexa Fluor 555-conjugated goat anti-mouse secondary antibody (1:1000 dilution, ThermoFisher, A32727). Cells were washed 3x with DPBS then coverslips were mounted using Prolong Diamond Antifade mountant (ThermoFisher, P36970). After 24 hours of curing, cells were imaged using a TissueFAXS SQL spinning disk confocal microscope with a 20X objective (TissueGnostics). Images were acquired using the “extended focus” setting, with one 1.5 μm step in both directions around the focus plane determined by autofocus in the DAPI channel.

### Assessment of proximity labeling by western blot

For each sample, 2e6 iBMDMs were seeded per well in a 24-well plate format. The following day, 10e6 of the appropriate beads (HRP- or BSA-beads) were prepared with 3x washes with D10 then added per well. iBMDM media was replaced with fresh D10 and HRP- or BSA-beads were added for a phagocytosis period of 30 minutes. To synchronize phagocytosis, extracellular beads were removed with 3x washes with warm DPBS, then warm media was re-added and phagosomes were allowed to mature for an additional 30 minutes at 37° C. AP (MedChem Express, HY-131442) or BP was spiked in to a final concentration of 500 μM at the indicated times prior to labeling. After 30 minutes of phagosome maturation, H_2_O_2_ was added to a final concentration of 1 mM with gentle agitation for 60 seconds, then the reaction was quenched by adding an equal volume of ice-cold 2X quenching buffer. Cells were washed 3x with ice-cold quenching buffer, detached and transferred to tubes, then pelleted and washed once more with ice-cold quenching buffer. Supernatant was removed and cell pellets were either stored at −80° C or proceeded to lysis immediately.

To ensure complete solubilization of membranes, cell pellets were briefly vortexed then lysed in a high-SDS lysis buffer (1% SDS in 50 mM Tris-HCl (pH 8.0) with 10 mM sodium azide, 10 mM sodium ascorbate, 5 mM trolox, 1 mM PMSF, and HALT protease inhibitors (ThermoFisher, 78429)). Lysates were boiled at 95° C for 5 minutes, then diluted with 4 volumes of 1.25x RIPA without SDS (50 mM Tris, 187.5 mM NaCl, 0.625% sodium deoxycholate, 1.25% Triton X-100) to achieve a final 1X RIPA lysis buffer with 0.2% SDS. Lysates were incubated at room temperature for 5 minutes, then clarified by centrifugation at 10,000xg for 10 minutes at 4° C. Lysates were quantified using the 660 nm Protein Assay with the Ionic Compatibility Detergent Reagent (ThermoFisher, 22663) and normalized to the same concentration with 1x RIPA buffer.

For click reactions, a 29x click master mix was prepared by first vortexing 14.5 mM CuSO4 with 29 mM BTTAA, then supplementing to the final concentrations of 72.5 mM sodium L-ascorbate and 2.9 mM biotin-PEG3-azide (Vector Labs, CCT-AZ104-25). The master mix was mixed by vortexing, then 2uL of master mix was added to 56 uL of lysates where indicated, and the reaction was carried out for 2 hours at 37° C with rotation. For “no click” conditions, 2 uL of water was added in place of the click master mix.Equal volumes of lysate were then loaded into a Bis-Tris gel (ThermoFisher, NW04120BOX) for electrophoresis, and transferred to a PVDF membrane. The PVDF membrane was stained with Ponceau S reagent (ThermoFisher, A40000279) for 5 minutes at room temperature, washed 2x with water for 60 seconds and imaged using a GelDoc (BioRad) for total protein normalization. The PVDF membrane was then blocked for 1 hour at room temperature with blocking buffer (Licor, 927-70001), washed 3x with tris-buffered saline + 0.1% tween-20 (TBS-T), and then stained for biotinylation with fluorescent streptavidin-800CW for 1 hour at room temperature (1:1000, Licor, 926-32230). The membrane was washed 3x with TBS-T, then once with TBS (no tween), prior to imaging with a Licor Odyssey CLx. Streptavidin and Ponceau S stain intensities were quantified using the “signal” quantification from ImageStudio (v5.2) with background subtraction, and the streptavidin intensity for each lane was normalized by dividing by the corresponding Ponceau S intensity for that lane. To compare intensities across replicates, each replicate was internally normalized by dividing each value from that blot by the intensity associated with the “BSA beads+BP+H_2_O_2_” condition for that replicate.

### Comparison of AP vs BP labeling on axenic beads

HRP- and BSA-beads underwent axenic labeling as described above. After 3x washes with quencher solution, beads were resuspended in 80 uL of quencher + 0.1% BSA. A 5x click master mix was prepared as described above and 20 uL were added to each sample as necessary and mixed with vortexing. The click reaction was carried out at 37° C for 1 hr with agitation, then washed 3x with 0.1% BSA and stained with streptavidin-AF647 and analyzed by flow cytometry as described above.

### Preparation of iBMDM samples for PhagoID proteomic analysis

35e6 IBMDMs were seeded into one 10cm dish ∼24 hrs prior to experiment for each condition. 250e6 beads were prepared per condition by washing 3x in D10. At time of experiment, fresh medium containing 250e6 beads, followed by 1 minute of agitation to evenly disperse beads, then 2 minutes with cells resting at room temperature to allow beads to settle with minimal internalization. Plates were then transferred to 37° C and allowed to internalize beads for 30 minutes. To synchronize phagocytosis, non-internalized beads were removed with 5x warm DPBS washes, then warm media was re-added and phagosomes were allowed to mature for an additional 30 minutes at 37° C. AP was spiked in to a final concentration of 500 μM 15 minutes prior to labeling. After 30 minutes of phagosome maturation, 100 mM H_2_O_2_ was prepared in DPBS and added to a final concentration of 1 mM for each dish, followed by 1 minute of gentle agitation, then rapidly removing the media and washing 3x with ice-cold quencher solution. Cells were then scraped in ice-cold quenching buffer and transferred to tubes, then pelleted and washed once more with ice-cold quenching buffer. Cells were pelleted again and supernatant was removed, then cell pellets were frozen at −80° C.

Cell pellets were thawed at room temp for 5 minutes, then vortexed briefly before adding 225 uL of 1% SDS lysis buffer (1% SDS, 50 mM Tris-HCl pH 8.0, 10 mM sodium ascorbate, 5 mM trolox, 1 mM PMSF, 1:1000 benzonase, 1:100 HALT protease inhibitor cocktail (ThermoFisher, 78429)) and incubating at RT for 3 minutes. Samples were boiled in 1% SDS lysis buffer at 95° C for 5 minutes, then 4 volumes of ice-cold RIPA (no SDS) were added and incubated for 5 minutes further at room temperature. Lysates were clarified by centrifugation at 10,000xg for 10 minutes at 4C and the clarified supernatant was quantified using the 660 nm Protein Assay with the Ionic Compatibility Detergent Reagent (ThermoFisher, 22663).

For the iBMDM proteomic experiment, all samples were normalized to 1 mg/mL in approximately 750 uL of 1% SDS lysis buffer. The click reaction was conducted with a 29x click master mix (resulting in final concentrations of 500 μM CuSO4, 1 mM BTTAA, 2.5 mM sodium L-ascorbate, 100 μM biotin-PEG3-azide) as described above at 37° C with rotation for 2 hours, then precipitated with 5 volumes of cold methanol overnight at −80° C. The next day, precipitated protein was pelleted by centrifugation at 8000xg for 3 minutes at 4C, and unreacted biotin-PEG3-azide was further washed away with 2x cold methanol washes. Lysate pellets were dried for 10 minutes, then resuspended in 100 uL of RIPA containing 2% SDS for 15 minutes with vigorous pipetting. 900 uL of RIPA containing no SDS was added to bring the total SDS to 0.2%, and lysates were transferred to fresh tubes prior to biotin enrichment by streptavidin beads. 75 uL of Pierce™ Streptavidin Magnetic Beads were washed 2x in RIPA then added to each sample and incubated at 4C with rotation overnight. Beads were then washed on ice with ice-cold buffers as follows: 2x washes with RIPA, once with 1M KCl, once with 0.1 M Na2CO3, once with 2 M Urea in 10 mM Tris-HCl (pH 8), and 2x with RIPA. Following the final RIPA wash, beads were transferred to fresh tubes and delivered to the Koch Institute Proteomics Core for further preparation for mass spectrometry. For the experiment testing the impact of PNGaseF digestion, samples were split at this point.

### Mass spectrometry

Samples underwent on-bead reduction, alkylation, and digestion. Proteins were reduced with 10mM dithiothreitol (Sigma) for 1h at 56° C and then alkylated with 20mM iodoacetamide (Sigma, I1149) for 1h at 25° C in the dark. Samples were incubated with PNGaseF for 2 h at 37° C. Proteins were then digested with modified trypsin (Promega, V5113) in 100mM ammonium bicarbonate, pH 8 at 25° C overnight. Trypsin activity was halted by addition of formic acid (99.9%, Sigma, 5.33002) to a final concentration of 5%. Peptides were desalted using Pierce Peptide Desalting Spin Columns (ThermoFisher, 89852) then vacuum centrifuged.

After digestion the samples were TMT labeled according to the manufacturer’s protocol (ThermoFisher, A44520). The dried tryptic peptides were resuspended in 100 mM TEAB, vortexed and then briefly centrifuged. 41 uL of anhydrous acetonitrile was added to each of the TMT label reagents, then vortex and briefly centrifuged. The TMT reagents were allowed to dissolve for 5 minutes. Then the 41 uL of the TMT reagents were added to each of the 100 uL of samples, vortexed and briefly centrifuged. The samples were allowed to incubate for 1 hour at room temperature, then 8 uL of 5% hydroxylamine was added to each sample and incubated for 15 minutes to quench the reaction. Equal amounts of each labeled sample were combined together and speed-vac to dryness. TMT labeled peptides were fractionated into 8 fractions for the iBMDM and hMDM PhagoID TMT 10plex experiments by Pierce High pH Reversed-Phase Peptide Fractionation Kit (ThermoFisher, 84868) per manufacturer’s instructions. Following fractionation, samples were speed-vacuumed to dryness.

Each fraction was separately resuspended in 0.2% formic acid and injected onto a column. The peptides were separated by reverse phase HPLC (Thermo Ultimate 3000) using a Thermo PepMap RSLC C18 column (2um tip, 75umx50cm, ThermoFisher, ES903) over a gradient before nano-electrospray using a Orbitrap Exploris 480 mass spectrometer (ThermoFisher). Solvent A was 0.1% formic acid in water and solvent B was 0.1% formic acid in acetonitrile. The gradient conditions were 1% B (0-10 min at 300nL/min) 1% B (10-15 min, 300 nL/min to 200 nL/min) 1-10% B (15-20 min, 200nL/min), 10-25% B (20-65 min, 200nL/min), 25-36% B (65-75 min, 200nL/min), 36-80% B (75-75.5 min, 200 nL/min), 80% B (75.5-80 min, 200nL/min), 80-1% B (80-80.1 min, 200nL/min), 1% B (80.1-100 min, 200nL/min).

The mass spectrometer was operated in a data-dependent mode. The parameters for the full scan MS were: resolution of 120,000 across 375-1600 m/z and maximum IT 25 ms. The full MS scan was followed by MS/MS for as many precursor ions in a two seconds cycle with a NCE of 36, dynamic exclusion of 30 s and resolution of 45,000.

Raw mass spectral data files (.raw) were searched using Sequest HT in Proteome Discoverer (ThermoFisher). Sequest search parameters were: 10 ppm mass tolerance for precursor ions; 0.05 Da for fragment ion mass tolerance; 2 missed cleavages of trypsin; fixed modification were carbamidomethylation of cysteine and TMT modification on the lysines and peptide N-termini; variable modifications were methionine oxidation, methionine loss at the N-terminus of the protein, acetylation of the N-terminus of the protein. As all samples included PNGaseF digestion, an additional variable modification of deamidation of asparagine because PNGaseF digestion of N-linked glycans results in the deamidation of the glycosylated asparagine. Data was searched against an Uniprot Mouse database (Proteome ID: UP000000589) and a contaminate database made in house.

### Mass spectrometry data processing for PhagoID experiments

We followed established analysis pipelines for analysis of the mass spectrometry data from phagosome proximity labeling mass spectrometry experiments^61,64^. Starting with the raw abundances of each protein as calculated by the Sequest HT search, a fold-change ratio was calculated for each protein by dividing the raw abundance in the HRP-bead sample by the raw abundance in the paired BSA-bead sample. This value was then log2-transformed. To normalize the data, the median log2-foldchange for each replicate was calculated among the proteins present within the FPlist (mouse or human Uniprot “mitochondrion matrix” proteome with proteins cross-listed as “endomembrane network” or “cytoplasm” removed), and this value was subtracted from the log2-foldchange for all proteins from that replicate. This centers the median false-positive protein at a log2-foldchange of 0 for each replicate. A moderated T-test was conducted using the limma package in R^122^ and p-values were corrected using the Benjamini-Hochberg protocol. For all mass spectrometry experiments, only proteins with an adjusted p-value of less than 0.05 were considered. For the iBMDM PhagoID experiment, the log2-foldchange cutoff was set such that 95% of proteins in the FP list (mouse-annotated Uniprot “mitochondrion matrix” proteome) had an average log2-foldchange below the threshold.

For the hMDM PhagoID experiment, the dataset was compared to a TP list set as the curated “murine core phagosome lumen proteome” derived from the iBMDM PhagoID experiment cross-referenced to previous existing datasets (Supplemental table 3). To establish a log2-foldchange cutoff, the TPR and were calculated for each log2-foldchange value by dividing the number of proteins in the TP list above the given cutoff by the total number of proteins above the given cutoff. The FPR was calculated for each cutoff as well by comparison to the FP list. The log2-foldchange was selected which maximized the specify, as measured by the delta between the TPR and FPR (TPR - FPR).

### Isolation of primary human CD14+ monocytes and human macrophage differentiation

CD14+ monocytes were isolated from deidentified buffy coats commercially acquired from Massachusetts General Hospital. Briefly, peripheral blood mononuclear cells (PBMCs) were isolated by density-based centrifugation and washing. Following isolation of PBMCs, CD14+ monocytes were isolated by immunomagnetic positive selection using a CD14+ selection kit (Stemcell Technologies, 17858). Isolated monocytes were differentiated into macrophages by culturing cells in macrophage differentiation media (Phenol-free RPMI (ThermoFisher, 11835055) supplemented with 10 mM HEPES (Corning, 25-060-Cl), 2 mM L-glutamine (Sigma, G-7513), and 10% heat-inactivated FBS (ThermoFisher, 10082147) and 25 ng/mL recombinant M-CSF (Biolegend, 574804)). Cells were cultured in ultra-low attachment T75 flasks (Corning, 3814) for the differentiation process to facilitate detachment and replating for downstream experiments.

### Preparation of hMDM samples for PhagoID proteomic analysis

hMDM samples for PhagoID were prepared identically to iBMDM samples as described above with the following modifications: For each hMDM sample, 25e6 CD14+ monocytes were isolated from healthy human donors as described above and were seeded onto low attachment T75 flasks and differentiated with M-CSF for 6 days. After 6 days of differentiation, cells were detached and hMDMs were plated onto 15cm dishes in macrophage media without M-CSF, and the experiment was carried out the following day. Following lysis and lysate quantification, all hMDM samples were normalized to 1 mg/mL in approximately 500 uL of 1% SDS lysis buffer before proceeding with the click reaction and further sample processing.

### GO and KEGG enrichment analysis

Analysis for the enrichment of GO and KEGG terms was performed using g:Profiler (version e111_eg58_p18_f463989d) with Benjamini-Hochberg multiple testing correction method and a significance threshold of 0.05^123^. The significant proteins (n=195 for iBMDM, n=123 for hMDM) were analyzed compared to a “custom background” consisting of all quantified proteins (n=2536 for iBMDM, n=1016 for hMDM).

### Isolation of iBMDM and hMDM phagosomes and analysis by PhagoFACS

Streptavidin M-280 Dynabeads (ThermoFisher) were washed 3x in DPBS then made fluorescent by conjugating with FITC-biotin at a ratio of 5 nmol FITC-biotin per 1 mg of beads (in a volume of 100 uL per 1 mg of beads) for 1 hour at room temperature with rotation. Beads were then washed 5x in the appropriate cell culture media before being added at a ratio of 10 beads per 1 cell to either iBMDMs or hMDMs, assuming 700 million beads per 1 mg of beads as reported by manufacturer. iBMDMs or hMDMs were allowed to phagocytose beads for an initial period of 30 minutes, then extracellular beads were washed away with 3x washes with warm DPBS. Warm cell culture media was re-added and cells were returned to the incubator for an additional 30 minutes of phagosome maturation.

Phagosomes were then isolated following the protocol reported by Fabrik and colleagues^44^. Briefly, cells were washed 1x with ice-cold DPBS, then scraped and pelleted on ice with DPBS. Importantly, samples were handled on ice or at 4° C in all following steps. The cell pellet was washed 2x with ice-cold DPBS. Cell pellet was resuspended in ice-cold homogenization buffer (water with 250 mM sucrose, 3 mM imidazole) for 5 minutes, then pelleted and resuspended in 1 mL of ice-cold homogenization buffer supplemented with HALT protease inhibitor cocktail (ThermoFisher). Samples were transferred to a 2 mL glass dounce homogenizer (Kimble, MilliporeSigma) and outer membranes were disrupted with 5 strokes with the tight-fitting pestle. Samples were then transferred into a fresh tube pre-coated with 1% BSA (diluted in DPBS). Samples were placed on a Dynamag-2 magnetic rack (on ice) and allowed to pellet for 90 seconds before removing supernatant and resuspending in fresh ice-cold 1% BSA. This wash was repeated 4x. To discriminate intact phagosomes prior to phagosome fixation and permeabilization, phagosomes were stained with a PE-conjugated anti-streptavidin antibody (1:100 dilution, Biolegend) in a 500 uL volume for 30 minutes on ice. Phagosomes were then washed 2x with ice-cold 1% BSA followed by 2x washes with ice-cold DPBS, then fixed with 4% PFA for 15 minutes at room temperature in 1 mL volume with rotation. Samples were washed 1x with ice-cold 1% BSA, then 3x with ice-cold DPBS to stop the fixation reaction. Phagosomes were then permeabilized in 100% methanol pre-chilled to −20° C for 30 minutes at 4° C in 1 mL volume with rotation, then washed 2x with ice-cold DPBS and 1x with ice-cold 1% BSA. To avoid non-specific binding of Fc receptors present on phagosomes, phagosomes were blocked by incubating in ice-cold 1% BSA + mouse Fc-block (1:100 dilution, Biolegend) + 0.3% Triton X100 on ice for 1 hour in 1 mL volume. Phagosomes were washed 2x with ice-cold 1% BSA, then stained with primary antibodies overnight at 4° C at the indicated dilution (Supplemental table 5). Phagosomes were washed 3x with ice-cold 1% BSA, then stained with fluorescent secondary antibodies for 1 hr on ice. Samples were washed 3x with ice-cold 1% BSA before analysis on an FACSymphony A5 flow cytometer (BD Life Sciences). All flow cytometry data was analyzed using FlowJo™ v10.8 Software (BD Life Sciences). Phagosomes were gated to only include intact phagosomes for analysis as described in Supplemental Figure 3.

## Data and materials availability

Our mass spectrometry data have been submitted to the PRIDE repository and will be made publicly available.

## Acknowledgments

We would like to thank members of the Bryson Lab for discussions and feedback. Additionally, we would like to acknowledge Alicia D’Souza, Daniel Maurer, Agnes Cheng, and Emerson Glassey for help in producing reagents and protocol optimization. We thank the Koch Institute’s Robert A. Swanson (1969) Biotechnology Center for technical support, specifically Rick Schiavoni and the staff of the Biopolymers & Proteomics Core.

## Supplemental Figure Captions

**Supplemental Figure 1: Alkyne phenol and biotin phenol yield equivalent labeling in axenic setting.**

A Representative flow cytometry distributions of beads labeled with biotin phenol (BP) or alkyne phenol (AP) followed by biotin-PEG3-azide click biotinylation where indicated.

B Quantitation and statistical analysis of (A) across independent replicates. (n=3 or 4)

**Supplemental Figure 2: Increased protein abundance upon digestion with PNGaseF is due to the detection of deamidated peptides.**

**Supplemental Figure 3: Quality control of iBMDM PhagoID experiment**

A Western blot of biotinylation across HRP-bead and BSA-bead samples from 5 independent replicates for iBMDM PhagoID mass spectrometry experiment.

B Heatmap displaying the Pearson’s correlation of protein quantification values from individual replicates from the iBMDM PhagoID mass spectrometry experiment.

**Supplemental Figure 4: PhagoFACS analysis distinguishes intact phagosomes from exposed beads and shows consistent heterogeneity of Cathepsins B and D.**

A Gating strategy for assessment of protein at intact phagosomes by PhagoFACS for both iBMDM and hMDM phagosomes. Gating hierarchy: Time Gate (not shown) > FITC singlets > Single phagosomes > Intact phagosomes.

B, C Consistency of Cathepsins B and D heterogeneity across multiple replicates in iBMDM (B) and hMDM (C) phagosomes.

**Supplemental Figure 5: Gene Ontology (GO) and KEGG enrichment analysis of 43 core phagosome lumen proteome proteins via g:Profiler.**

**Supplemental Figure 6: Normalized mass spectrometry abundance of 27 novel proteins detected at the phagosome by PhagoID.**

**Supplemental Figure 7: Quality control of hMDM PhagoID experiment**

A Western blot of biotinylation across HRP-bead and BSA-bead samples from 5 independent donors for hMDM PhagoID mass spectrometry experiment.

B Heatmap displaying the Pearson’s correlation of protein quantification values from individual donors from the hMDM PhagoID mass spectrometry experiment.

C Receiver-operator-curve (ROC) analysis of hMDM PhagoID compared to the true positive list (core phagosome lumen proteome) and the false positive list (Uniprot mitochondrial matrix proteome with proteins cross-listed as “endomembrane network” or “cytoplasm” removed). TPR = true positive rate, FPR = false positive rate.

D Selection of the optimal fold-change threshold based on the maximized TPR - FPR value.

**Supplemental Figure 8: Data supporting differences between iBMDM and hMDM phagosome proteome**

A Normalized mass spectrometry abundance of proteins detected as being differentially abundant at the iBMDM and hMDM phagosome lumen by PhagoID.

B Western blots of three proteins with cross-reactive antibodies (top) with total protein staining by Ponceau S (bottom).

C Normalized quantitation of western blot intensity from (B). (n=1)

**Supplemental Tables**

**Supplemental Table 1: iBMDM PhagoID proteomics data.**

**Supplemental Table 2: g:Profiler enrichment analysis of iBMDM and hMDM PhagoID datasets.**

**Supplemental Table 3: Comparative analysis between PhagoID and previous phagosome proteome studies.**

**Supplemental Table 4: hMDM PhagoID proteomics data. Supplemental Table 5: Description of antibodies used in this study.**

