## Supplementary figures and images for "Proximity labeling defines the phagosome lumen proteome of murine and primary human macrophages"

### Supplemental Figure 1

**Supplemental Figure 1**

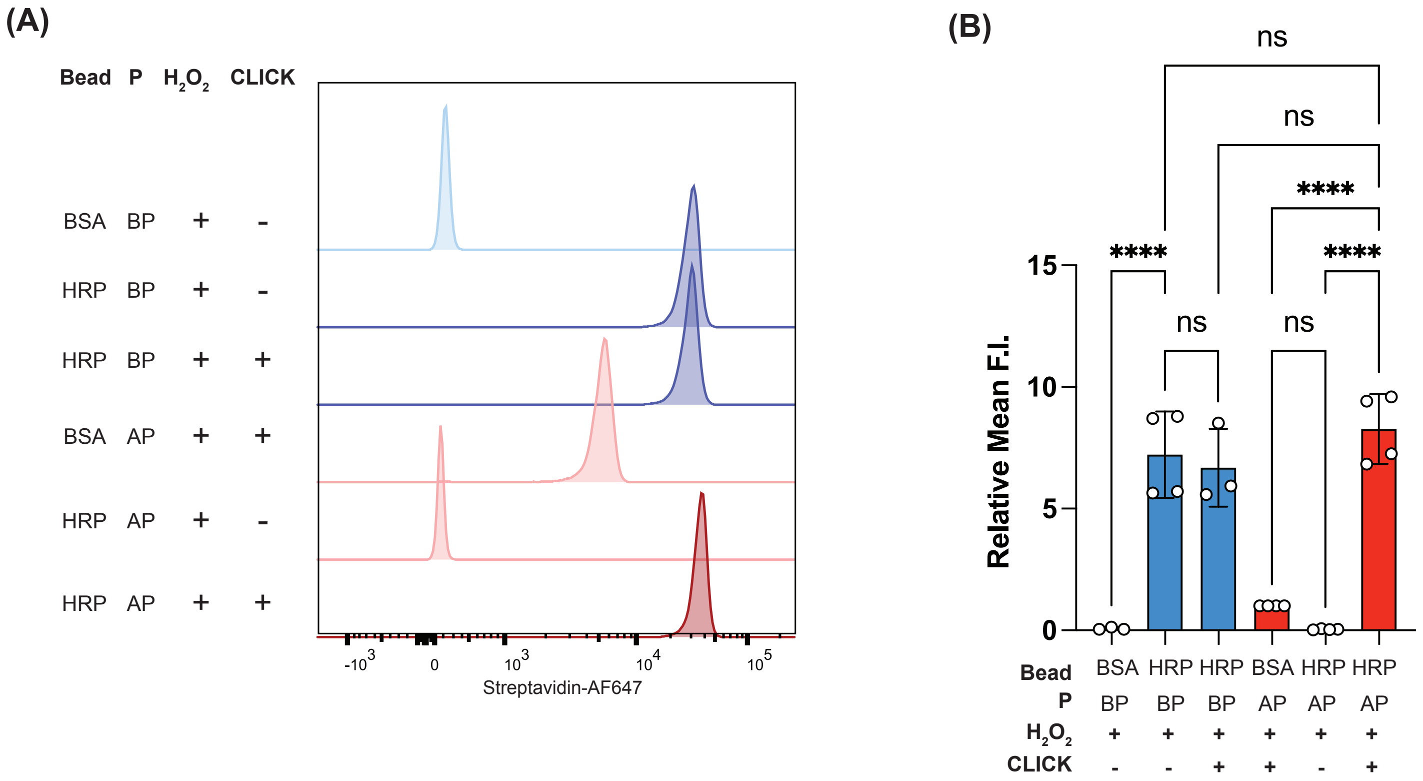

### Supplemental Figure 2

Supplemental Figure 2

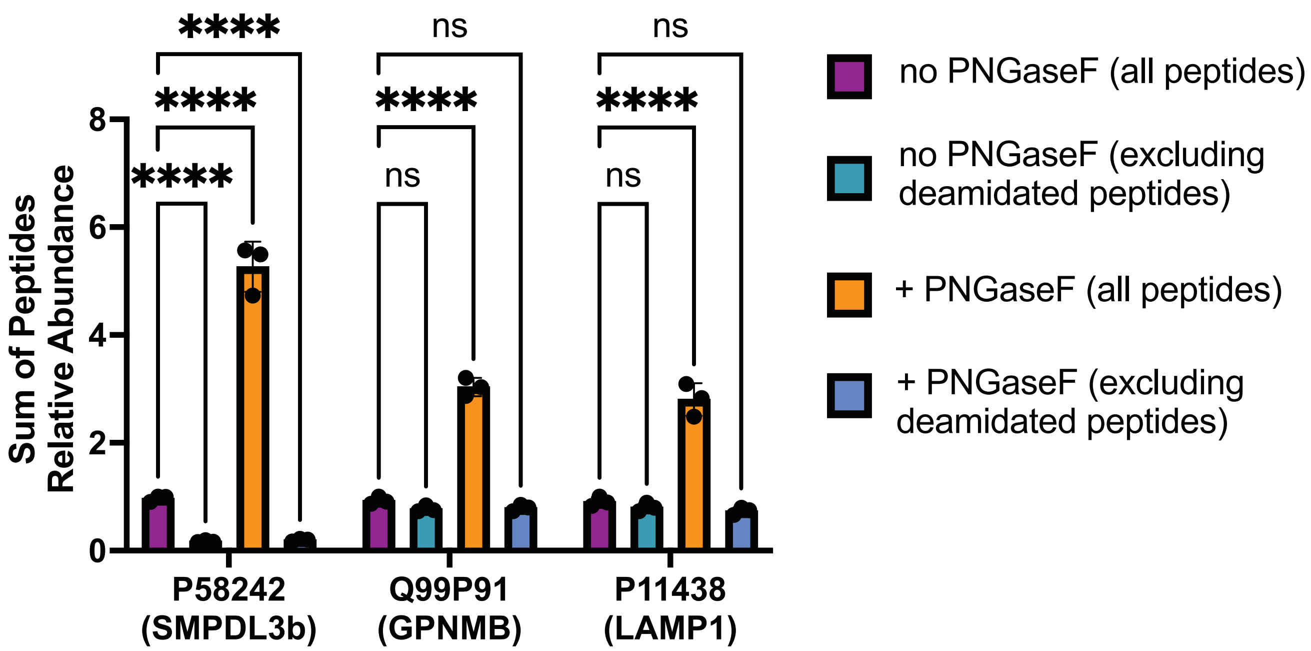

### Supplemental Figure 3

#### Supplemental Figure 3

(A)

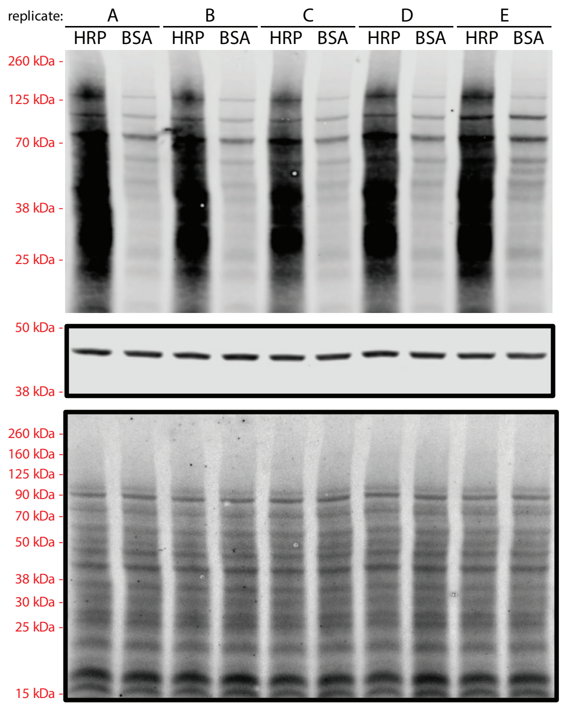


(B)

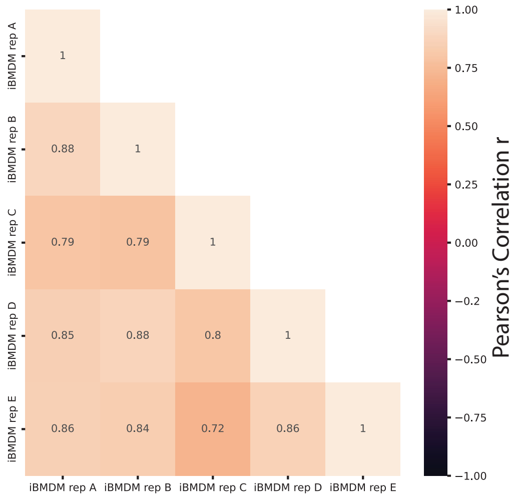

### Supplemental Figure 4

**Supplemental Figure 4**

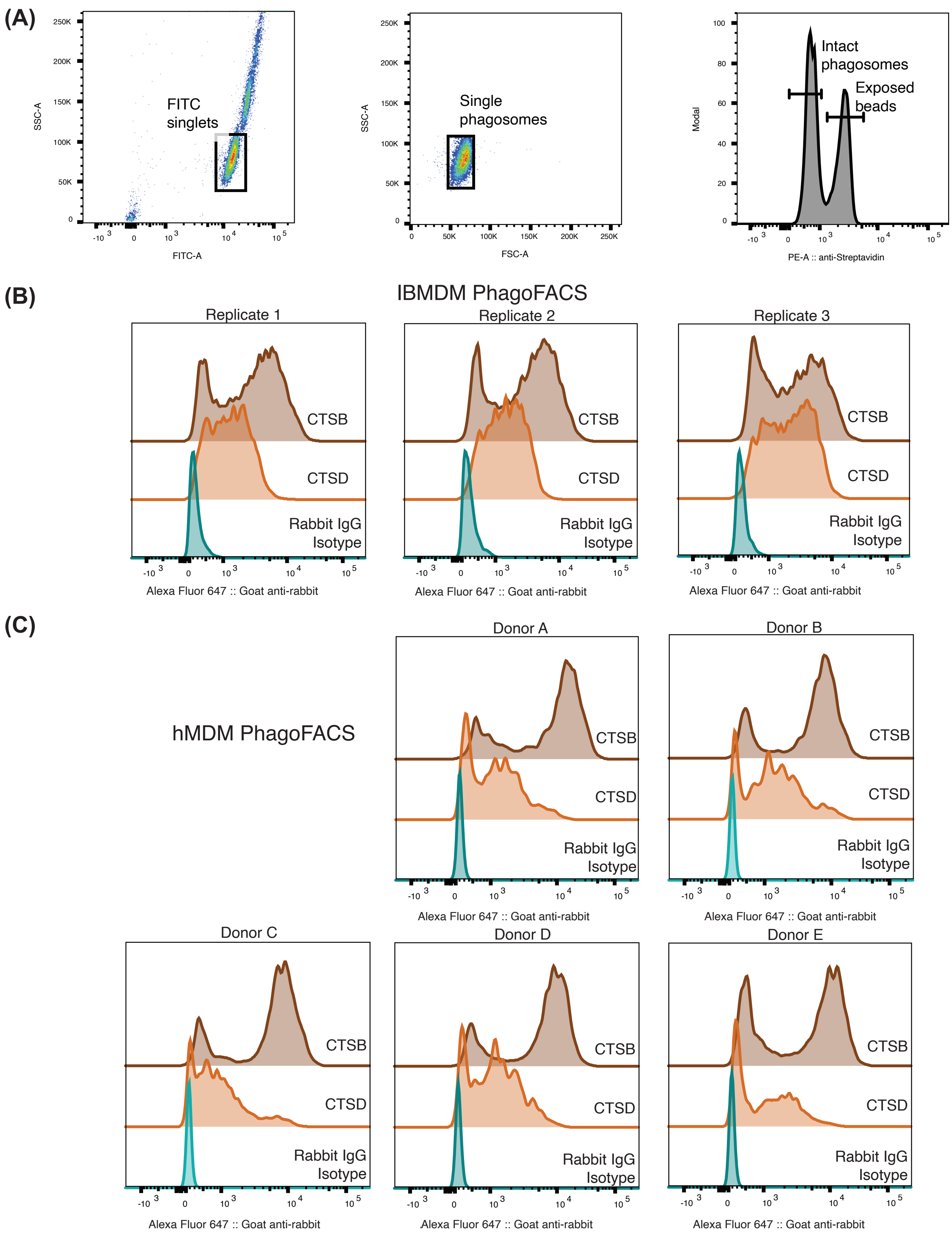

### Supplemental Figure 6

**Supplemental Figure 6**

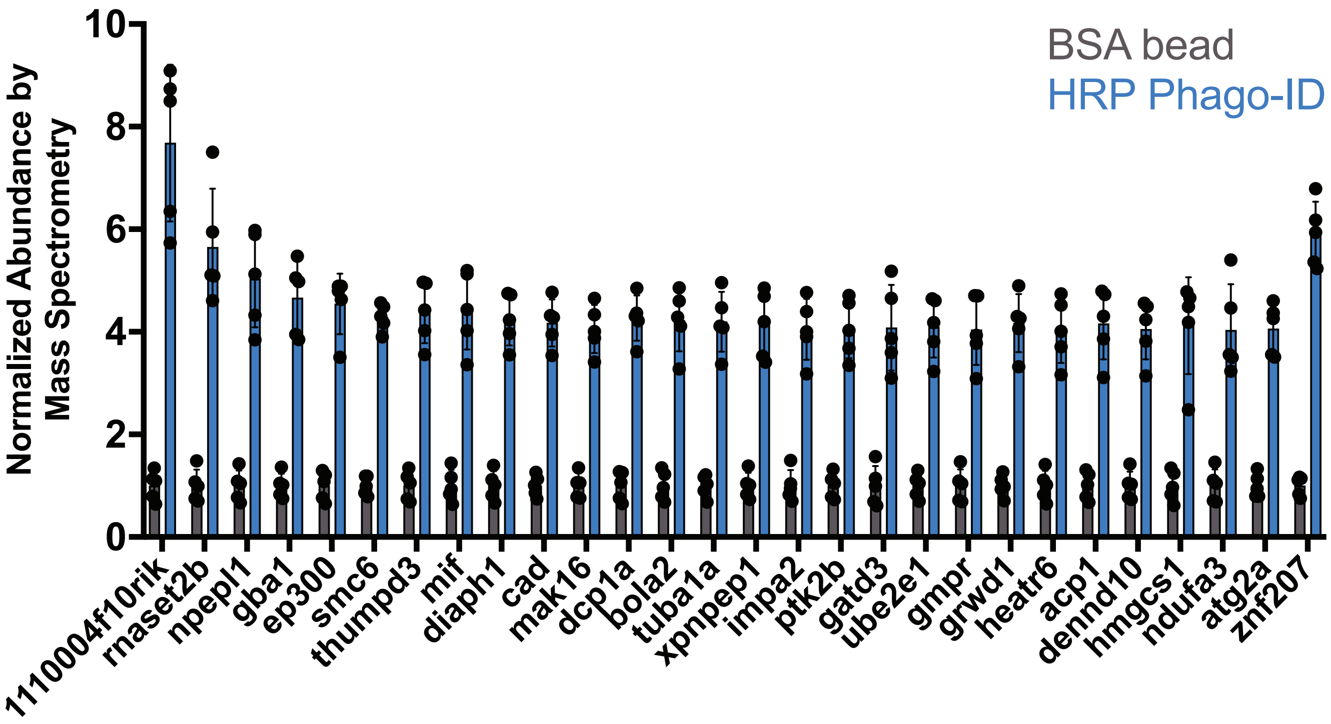

### Supplemental Figure 7

##### Supplemental Figure 7

(A)

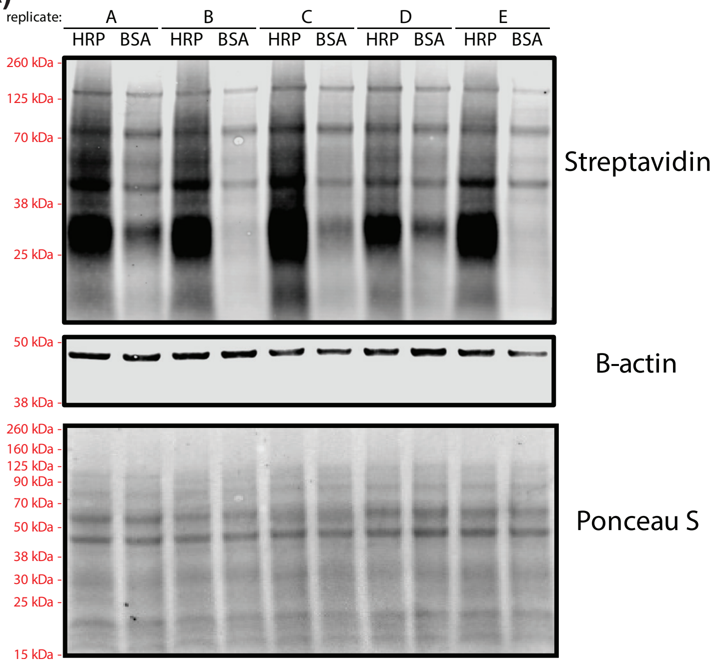


(B)

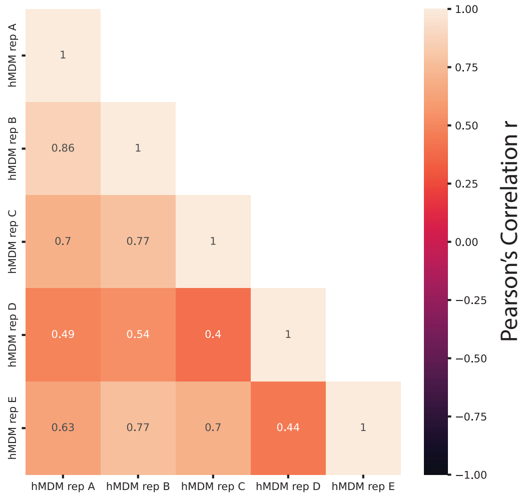


(C)

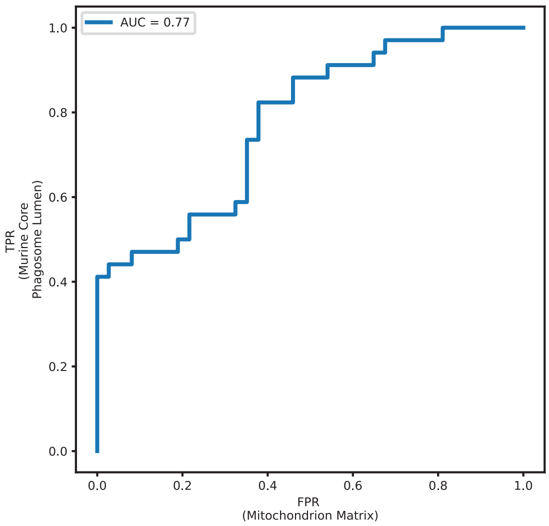


(D)

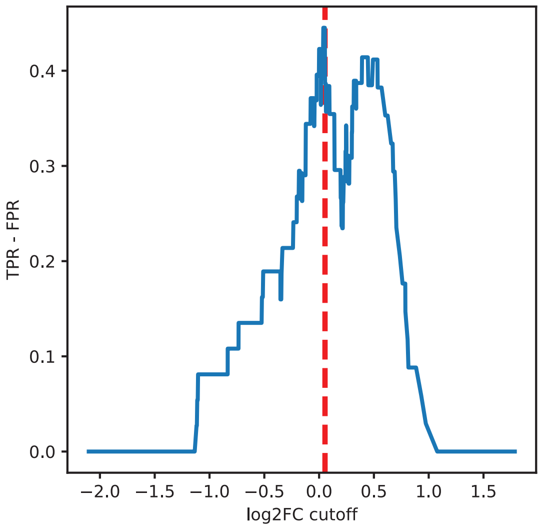

### Supplemental Figure 8

Supplemental Figure 8

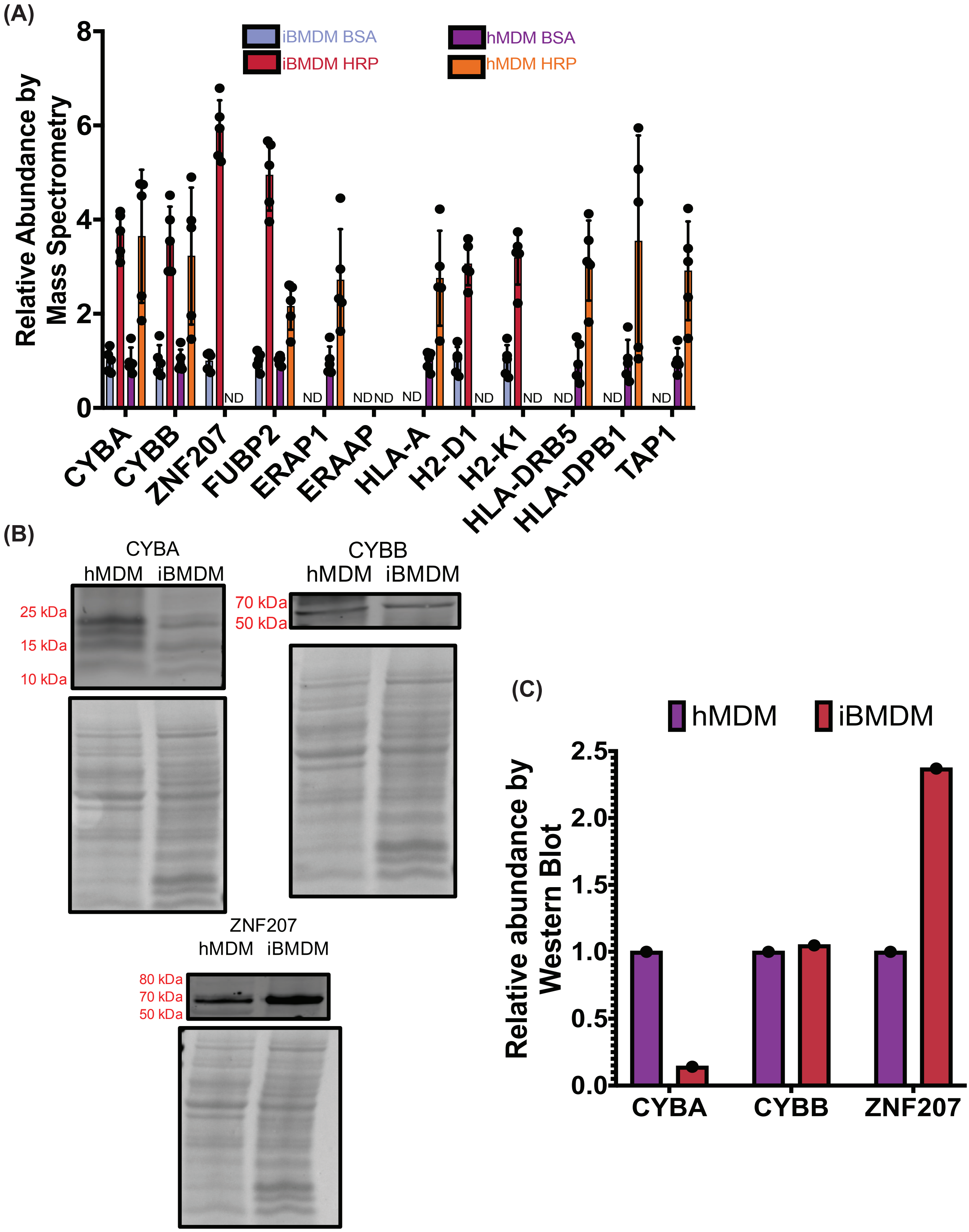
