## Supplemental Figure 5 for "Proximity labeling defines the phagosome lumen proteome of murine and primary human macrophages"

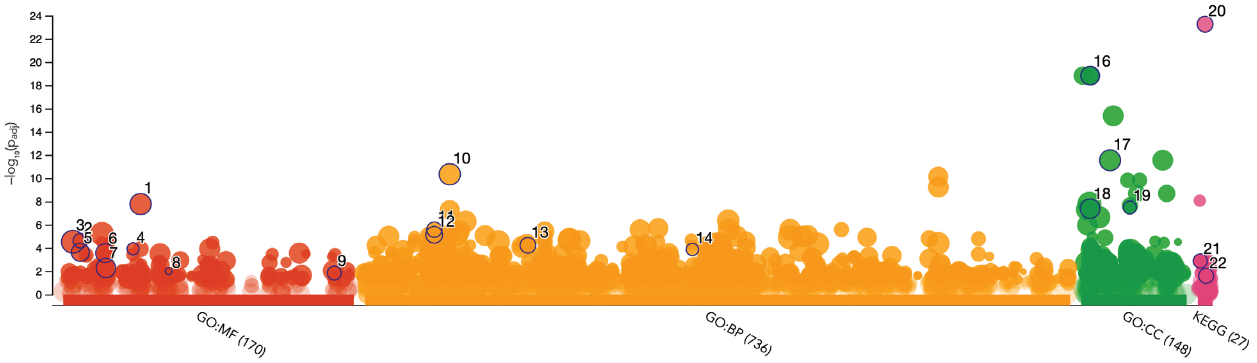


| ID | Source | Term ID | Term Name | $p_{adj}$ (query_1) |
|---|---|---|---|---|
| 1 | GO:MF | GO:0016787 | hydrolase activity | $1.683\times10^{-8}$ |
| 2 | GO:MF | GO:0004197 | cysteine-type endopeptidase activity | $2.401\times10^{-5}$ |
| 3 | GO:MF | GO:0003824 | catalytic activity | $2.943\times10^{-5}$ |
| 4 | GO:MF | GO:0015929 | hexosaminidase activity | $1.189\times10^{-4}$ |
| 5 | GO:MF | GO:0004175 | endopeptidase activity | $2.364\times10^{-4}$ |
| 6 | GO:MF | GO:0008233 | peptidase activity | $2.770\times10^{-4}$ |
| 7 | GO:MF | GO:0008289 | lipid binding | $5.324\times10^{-3}$ |
| 8 | GO:MF | GO:0030882 | lipid antigen binding | $1.002\times10^{-2}$ |
| 9 | GO:MF | GO:0140313 | molecular sequestering activity | $1.447\times10^{-2}$ |
| 10 | GO:BP | GO:0009056 | catabolic process | $4.555\times10^{-11}$ |
| 11 | GO:BP | GO:0007040 | lysosome organization | $2.644\times10^{-6}$ |
| 12 | GO:BP | GO:0007033 | vacuole organization | $7.377\times10^{-6}$ |
| 13 | GO:BP | GO:0019882 | antigen processing and presentation | $5.887\times10^{-5}$ |
| 14 | GO:BP | GO:0046514 | ceramide catabolic process | $1.349\times10^{-4}$ |
| 15 | GO:CC | GO:0005773 | vacuole | $1.501\times10^{-19}$ |
| 16 | GO:CC | GO:0005764 | lysosome | $1.501\times10^{-19}$ |
| 17 | GO:CC | GO:0031410 | cytoplasmic vesicle | $2.864\times10^{-12}$ |
| 18 | GO:CC | GO:0005768 | endosome | $4.168\times10^{-8}$ |
| 19 | GO:CC | GO:0043202 | lysosomal lumen | $3.290\times10^{-8}$ |
| 20 | KEGG | KEGG:04142 | Lysosome | $5.549\times10^{-24}$ |
| 21 | KEGG | KEGG:00600 | Sphingolipid metabolism | $1.328\times10^{-3}$ |
| 22 | KEGG | KEGG:04612 | Antigen processing and presentation | $2.518\times10^{-2}$ |

**version** e111_eg58_p18_f463989d
**date** 8/28/2024, 12:15:06 PM
**organism** mmusculus

g:Profiler
